# Novel Tissue Mechanics-Guided Cellular Flows Enable the Evolution of Feather Follicles

**DOI:** 10.1101/2025.10.01.679696

**Authors:** Hans I-Chen Harn, Ting-Xin Jiang, Chih-Han Huang, Wen-Tau Juan, Tzu-Yu Liu, Tsao-Chi Chuang, Wan-Chi Liao, Yingxiao Wang, Ji Li, Cornelis J. Weijer, Ping Wu, Chin-Lin Guo, Cheng-Ming Chuong

**Affiliations:** Department of Pathology, Keck School of Medicine, University of Southern California, Los Angeles, CA, USA; Ostrow School of Dentistry, University of Southern California, Los Angeles, California, USA; Division of Cardiovascular Surgery, Department of Surgery, Tri-Service General Hospital, National Defense Medical Center, Taipei, Taiwan; Department of Biomedical Imaging and Radiological Science, China Medical University, Taichung, Taiwan; Department of Medical Research, China Medical University Hospital, Taichung, Taiwan; Department of Life Sciences, National Cheng Kung University, Tainan, Taiwan; Department of Bioengineering & Institute of Engineering in Medicine, University of California, San Diego, La Jolla, California, USA; Alfred E. Mann Department of Biomedical Engineering, University of Southern California, Los Angeles, California, USA; Department of Dermatology, Xiangya Hospital, Central South University, Changsha, China; School of Life Sciences, University of Dundee, Dundee, United Kingdom; Institute of Physics, Academia Sinica, Taipei, Taiwan. 128 Sec 2 Academia Rd, Nankang District, Taipei, Taiwan

## Abstract

Complex tissue architecture is achieved through multiple rounds of morphological transitions. Here, we analyzed cellular flows and tissue mechanics during avian skin development. We showed how novel cellular flows initiate chemo-mechanical circuits that guide epithelial protrusion, folding, invagination, and spatial cell fate specification. In the initial feather bud formation, stiff dermal condensates protrude vertically out of the locally softened epithelial sheet. As the bud elongates, it stretches the epithelial cells at the base, which mechanically activates YAP and causes the epithelial sheet to fold downward and form a stiff cylindrical wall that invaginates into the skin. This stiff epithelial tongue is essential to the compaction and formation of the tightly packed dermal papillae. These topological transformational events are mechanically interconnected, and the completion of the previous circuit initializes the next one. On the contrary, during scale development, its rigid epithelial sheet restricts dermal cell flows, preventing other further topological transformation. We generated a topological transformation model to show how the process enables the novel evolution of feather follicles.

## Main text

The integument forms the boundary between the organism and its environment and evolves according to essential functioning. In early chordates, such as amphioxus, integuments are smooth and lack appendages. In fish, scales begin to form through epithelial placode formation. More complex skin appendages gradually emerge, helping vertebrates adapt to different ecological niches (Dhouailly, 2023; Wu *et al*, 2004). Various types of skin scales, glands, and spines are generated through simple topological transforming processes such as epithelial folding, invagination, protrusion, and branching. The more complex structures are follicles, found in feathers, hair, and reptile teeth (Fig. S1A). While these follicles evolved independently, they share characteristics such as having stem cells and a dermal niche, and an architecture that allows follicles to undergo cyclic renewal throughout the organism’s lifetime (Fig. S1B).

Among these, feather follicles are the most complex. The filament cylinder contains progenitor cells in the proximal end(Yue *et al*, 2005) that undergo branching morphogenesis toward the distal end (Li *et al*, 2017; Yu *et al*, 2002). Dermal papilla not only has inducing ability for epidermis but also undergo cyclic renewal itself (Wu *et al*, 2021b). At various ages, mature feathers exhibit different phenotypes (Chen *et al*, 2015; Chen *et al*, 2024). This is made possible because feathers have follicles that house stem cells (Yue *et al*., 2005) and can undergo cyclic regeneration.

During the evolution of feathers, the ability to molt and regenerate is a pre-requisite for the ability to generate diverse feather forms. While the evolutionary development (evo-devo) of feather forms has been extensively studied, tracking back to feathered dinosaurs (Li *et al*., 2017; Prum, 2005; Xu *et al*, 2014), few studies have been focused on the evolution of feather follicles. Based on fossil findings, it has been suggested that the two different feather forms in oviraptorisaur Similicaudipteryx (Prum, 2010; Xu *et al*, 2010) may represent the primary feather transition process we observed in today’s birds (Chen *et al*., 2024). By analyzing flight feather fossils, it also has been suggested that feathers are able to molt sequentially in Microraptor (Kiat *et al*, 2020) and Mesozoic birds (Wang *et al*, 2024). These processes require molting and regeneration of new feathers from the follicle, suggesting the existence of ancient feather follicles in feathered dinosaurs. On the other hand, diffusible morphogen such as Shh, Wnt, BMP, Eda are present in the reptile scales, other planar integument as well as in feather morphogenesis (Chen *et al*., 2015; Di-Poi & Milinkovitch, 2016). With similar molecular tools available, what event drives the planar epithelia to be transformed into a complex architecture such as the feather follicle (Fig. S1)?

Here we take a biophysical approach to study how a feather follicle is built and evolved, aiming to understand key biomechanical - biochemical crosstalk. The morphogenetic process can be understood through various levels of symmetry-breaking events (Prum, 2005) that involve cell migration and tissue folding, with cellular forces and migration serving as crucial drivers that initiate and execute these processes, leading to architectures ranging from simple to complex (Collinet & Lecuit, 2021; Tozluoglu *et al*, 2019). Due to the viscoelastic nature of biological systems, sculpting biological structures entails transitioning constituent materials between liquid to solid-like states (Lenne & Trivedi, 2022), as reflected in changes of cellular movements and mechanical properties. The capacity to increase or decrease local tissue stiffness and fluidity, in turn, creates the stiffness difference that allow cells to collectively migrate or “flow” accordingly, leading to the sculpture of the organ in the making. For example, in zebrafish development, the tissue undergoes jamming and unjamming of cell flows and solid to fluid transition to achieve the unidirectional axis elongation (Mongera *et al*, 2018). During Drosophila wing imaginal disc development, multiple epithelial invaginations and buckling are required to generate localized and anisotropic forces for the epithelial cells to initiate symmetry breaking and tissue folding (Bailles *et al*, 2019; Tozluoglu *et al*., 2019). However, the chemo-mechanical spatial regulation of morphogenesis in bi-layered composite materials involves continual dynamic interactions between the epithelia and mesenchyme, and how such dynamics lead to the building of complex ectodermal organ such as the feather remains to be investigated.

In different stages of avian skin development - feather bud formation, follicle invagination and dermal papillae formation - epidermal and dermal cells migrate synergistically, transitioning from a simple 2D placodal plane into a 3D follicle architecture that encloses different spatially specified cell fates (Chuong *et al*, 2000). How does epithelial tissue fold affect dermal cell flows and fate specification? How do chemo-mechanical factors control anisotropic tissue expansion, from simple to complex multi-dimensional tissue structures? In this study, we use chicken and transgenic quail skin explant models to characterize cellular flows, tissue mechanics, chemo-mechanical coupling molecules, and simulate the interactions through modeling during different stages of feather development. We selected exemplary key molecules and identified their role in not only molecular signaling but also in initiating key mechanical events that facilitate sequential stages of morphogenesis. Using these cellular flow, spatial mechanical and morphogenetic molecular findings, we generated a mechano-chemical coupling model for follicle formation that describes the initiating and extending of epidermal invagination around the dermal condensate (DC) and the eventual establishment of dermal papillae (DP). This model distills the essential parameters for such topological transformation. We test the prediction of this model in scutate scale formation and the induction of feather follicles from these scales (Lai *et al*, 2018; Wu *et al*, 2018b). This work highlights the formation of follicles resulting from an evolutionary novel morphogenetic process. New cell flows driven by mechanics allow the stem cell-based follicle to be built.

### Results

#### Stiff dermal cell condensate vertically protrudes locally softened epithelial sheet during bud formation

To characterize the cell flow dynamics, we recorded confocal time-lapse videos used either H2B-mOrange labelled chicken embryo or mCherry-H2B labelled transgenic Japanese quail explants. During early feather bud formation (E7-E8), dermal condensations (DC) form via chemo-mechanically sensitive cell proliferation and migration, such as those mediated by Fgf (Song *et al*, 1996; Widelitz *et al*, 1996), Dermal condensation formation is characterized by dermal cells that first migrate horizontally, which later protrude out to form the initial feather bud (Fig 1A). The bud shown is in the short bud stage, which later elongates to form the long bud (Li *et al*, 2013; Li *et al*., 2017). Here we focus on the formation of the short bud. Cell tracking analysis demonstrating the changes of cellular flow directions during different stages of feather bud formation (Fig 1B, Supplementary Video 1). In the first 6h of culturing, most of the dermal cell movements occur in the horizontal (xy) plane parallel to the epidermal layer, forming the DC, which is accompanied by a decrease in the distance between neighbouring cells (SI Fig 2B). Surprisingly, between E7 6-18h, the DC cells begin to move vertically (z) in a direction perpendicular to the epidermal layer, forming the protrusion of the early feather bud (Fig 1B).

**Fig. 1.**
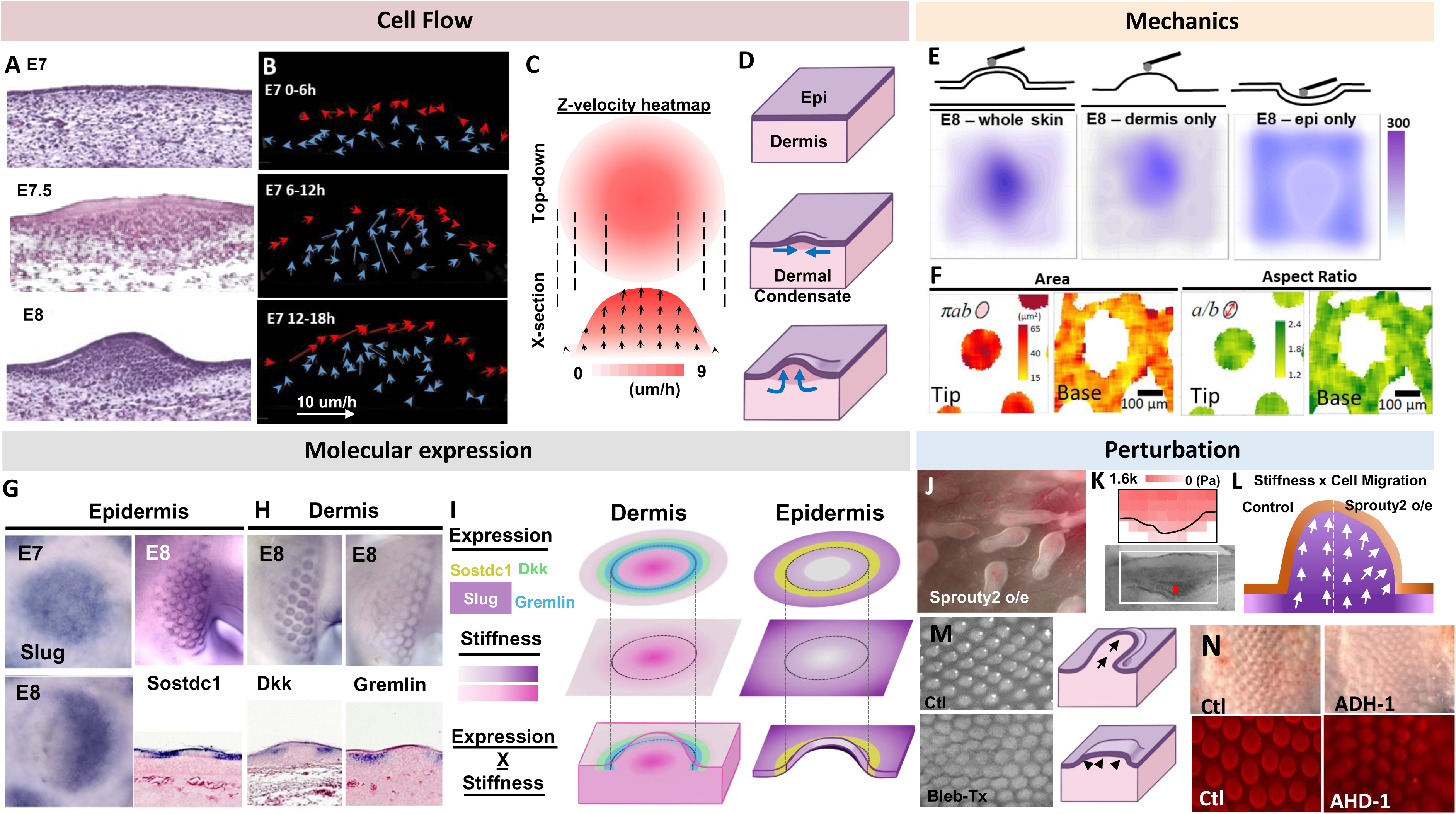
Feather bud protrusion: Vertical cellular flows and the formation of bud boundary. A. H&E images of chicken skin development from E7 to E8. B. Cell tracking analysis of E7 chicken explant culture for 18h reveals epidermal (red arrows) dermal cell (blue arrows) migration pattern changed from horizontal to vertical during bud protrusion. The length and direction of the arrows indicate the velocity of the cells tracked. C. Z-velocity heatmap showing cells at the center and apical region of feather bud migrate distally faster than those at the periphery. The direction and length of the arrows reflect the average velocity of cells at that specific region. D. Illustration of cellular flows at different stages of early feather bud formation. E. Landscape of tissue stiffness measured by AFM. Shown here are E8 whole skin, or dermis, epidermis alone. The data suggest stiffness mainly come from dermis at this stage. F. QMorF analysis showing the force distribution reflected by the epidermal cells in terms of cell area and aspect ratio during feather bud protrusion. G-H. In situ hybridization of selective molecules. G. Slug and WISE/Sostdc1 in E7 and E8 chicken skin epidermis. H, DKK1 and Gremlin in E8 chicken skin dermis. Slug is on the placode epithelium. Remarkably Sosdc1, DKK and Gremlin show ring staining pattern of different radius in whole mount. Section staining show epidermal staining for SosDc1, and Dkk (inside) and Gremlin (outside) flank bud boundary. I. Illustration of spatial distribution of epidermal (purple) and dermal (pink) tissue stiffness and molecular expression in an early feather bud. Darker color indicates higher stiffness. J. Sprouty2 overexpression in the epidermis cause bud to shorten and expand distally. Representative photo. K. Characterization of Sprouty perturbed buds. AFM stiffness heatmap and bright view photo of a bulging feather bud on one side of a locally Sprouty2-overexpressing feather filament. L. Stiffness and cell migration illustration. Epidermis softening through Sprouty2 overexpression on one side of a bud. On the perturbed side, dermal cell migration is directed more laterally and hence leading to a bulging feather bud. M. Blebbistatin treatment (Bleb-Tx) lead to shortened and widened feather buds in E9 explants cultured for 48h. This is correlated with softened dermal cell rigidity and abrogated vertical cell migration. N. Bright field and PI-stained photos of E8+48h chicken skin explants with NCAD inhibitor (ADH-1) show smaller and shorter feather buds.

This is accompanied by the increase in the average distance to the nearest cells (SI Fig 2B, late protrusion). The z-velocity heatmap during the first 16h recording shows that the cells at the center of the feather bud migrate faster than those at the periphery, and cells at the apical side migrate faster than those at the basal side (Fig 1C). The illustration shows dermal cells undergo a switch in movement from the horizontal to vertical cellular direction during feather bud protrusion (Fig 1D). With changes of dermal cell movement in both direction and speed, this new cellular flow implies a switch to a more fluid behaviour along the Z axis (Mongera *et al*., 2018) (SI Fig 2B).

How do these dermal cells change their migration direction to move upward? Considering that the FGF-stimulated DC cells exhibit high motility and are confined by the epidermis, one likely mechanism to initiate their vertical upward motion is to have a mechanically weak point at the epidermis above the DC. We used atomic force microscopy (AFM) and an image-based quantitative morphology field (QMorF) measurement (Wu *et al*, 2021a) to investigate the spatial mechanical dynamics and the associated cellular morphing of epidermis and dermis in E8 feather bud in contrast to surrounding non-bud regions. The stiffness heatmaps demonstrate that the whole skin feather bud is more rigid (0.273 ± 0.031 kPa) than the surrounding peri-bud region (0.187 ± 0.019 kPa, Fig 1E, left). We then separate the epidermis and dermis to examine their contributions to tissue rigidity separately. The dermis shows a similar stiffness difference, as well as an overall decrease in stiffness (bud: 0.239 ± 0.022 kPa, inter-bud: 0.178 ± 0.019 kPa). Interestingly, the peeled epidermis shows an opposite stiffness map. The bud region is softer (0.189 ± 0.017 kPa), while the inter-bud region is stiffer (0.209 ± 0.021 kPa, Fig 2E) than that of the dermis (SI Fig 2C). We also found that the regional stiffness of the bud region also increases during dermal condensation and early feather bud protrusion (SI Fig 2D). These findings suggest that the vertical movement of DC cells during feather bud protrusion is guided, at least partially, by this locally softened epidermis. The finding that DC cells had a stiff base to prevent them from invaginating downwards into the dermis is consistent with this observation (Yang *et al*, 2023). Thus, the decreased mechanical resistance at the epidermal placode allows dermal cells of DC to “flow out” of the skin surface.

**Fig. 2.**
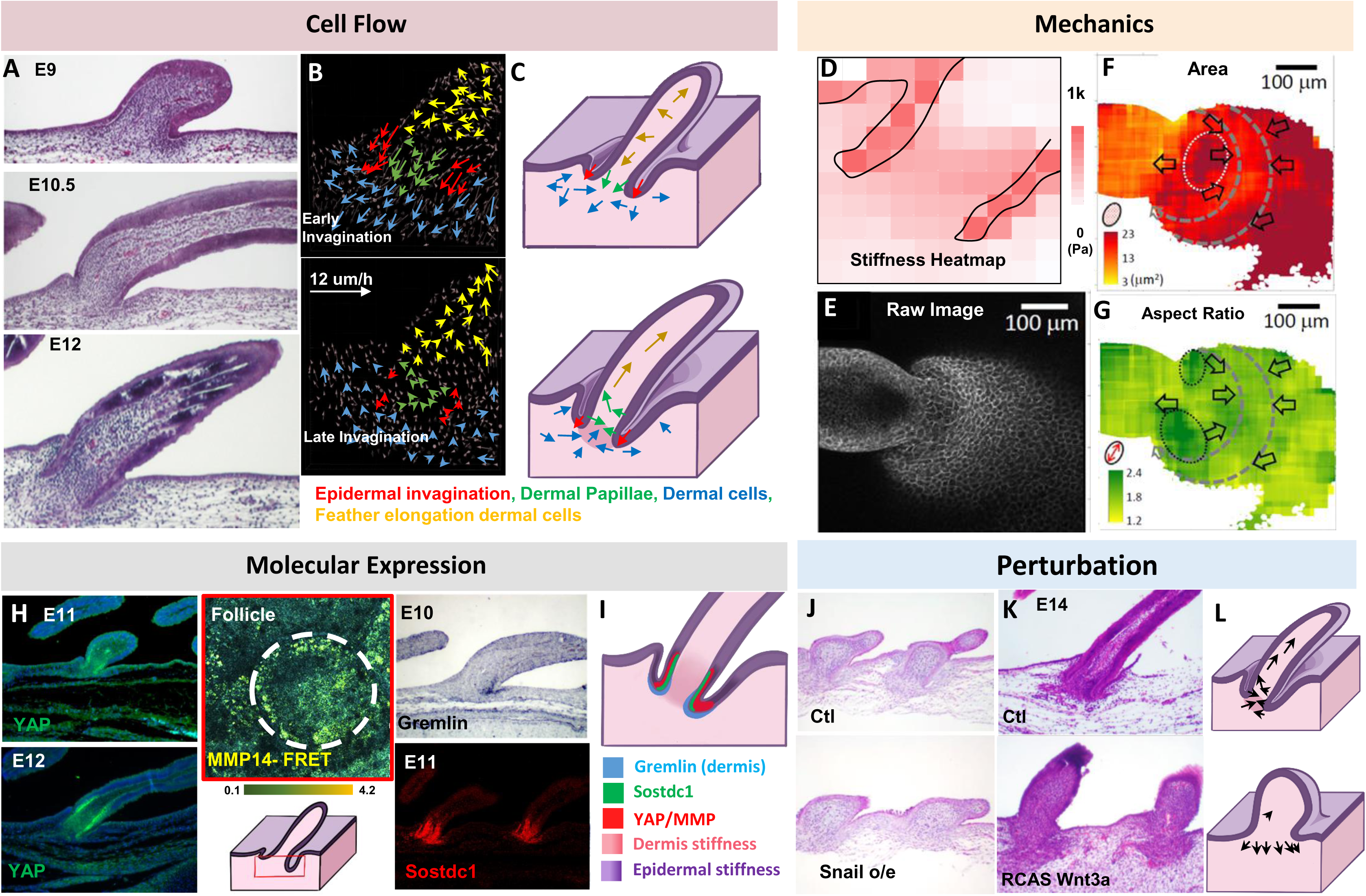
Feather follicle invagination: Invaginating epidermal cells compartmentalize and create differential dermal cell flows. A. H&E images of feather follicle invagination at E9, E10.5 and E12. B. Cell track analysis of E10+24h showing different cellular flows: (red arrows) epidermal invaginating cells, (green arrows) dermal papillae cells, (blue arrows) dermal cells, and (yellow arrows) feather elongation dermal cells. C. Illustration of cellular flows during early (upper) and late (lower) follicle invagination. The red arrows indicate invaginating epidermis that form as physical barriers and creates differential cell flows in the dermis, DP and feather bud. D. Stiffness heatmap of E10 follicle invagination in sagittal section of a follicle. Measured using AFM. Black lines demarcate epidermis. E. Whole mount E10 feather bud with invagination stained with E-cadherin. F. QMorF analysis of panel E shows force distribution reflected by the epidermal cells during feather follicle invagination. Arrows indicate local force directions estimated by the gradient of cell area, considering each cell as an embedded local force gauge. G. QMorF analysis. Aspect ratio of E10 epidermal cells in and around a feather bud and invagination site. H. In situ hybridization of Gremlin, RNAscope of Sostdc1, immunohistochemistry of YAP, and MMP14-FRET activity images in E10-E12 chicken skin during follicle invagination. Note the specific expression of YAP in proximal bud epidermis, some cells show cytoplasmic stain, some show nuclear stain. Sostdc1 in proximal bud dermis. I. Schematic summary showing Gremlin, Sostdc1, YAP and MMP expression and spatial tissue stiffness of an E12 follicle. Darker color indicates higher tissue stiffness. J-L Perturbation of follicle formation by interrupting mechano-chemical coupling between epidermis and dermis. J. H&E photos of Ctl and Snail overexpression in the epidermis inhibited epidermal invagination in E7 reconstituted skin for 7 days. K. H&E photos of E14 control and Wnt3a overexpression abrogated epidermal invagination, leading to dispersed dermal condensate cells, shortened feather buds and no follicle wall invagination. L. Illustration of abrogated epidermal invagination via softening epidermis causes dispersal of dermal condensate cells, and in turns leads to shortening and widening of feather buds.

For force balance to occur, the upward vertical movement of dermal cells must be accompanied by reciprocal downward vertical forces exerted at the epidermal layer. We wonder what reciprocal physical-morphogenic profile that the epidermal cells exhibit during bud protrusion. We use cell shape changes (by E cadherin staining) and QMorF method (Chang *et al*, 2019; Wu *et al*., 2021a) to analyze the changes in cell shape dynamics at the skin surface. We observed that during early bud formation, the angle and aspect ratio of epidermal cells are converging and stretched towards the feather bud (Fig 1F, SI Fig 3A-C). The initial analyses focused on cross-sections at different levels from the top (z=54 μm) to the base (z=0) of a developing feather bud at the E9 stage (Fig. 1F). The raw fluorescence image for each optical section is displayed in (SI Fig. 3C). QMorF distribution heatmaps of the coarse-grain averaged cross-sectional morphological characteristics (SI Fig 3B) including area (πab), aspect ratio (a/b), and orientation (θ) were illustrated in SI Fig 3D, E, and F, respectively. SI Fig 3D revealed that the average area decreases from the top to the base of the feather bud, suggesting that cells at the top experience greater “stretching” forces than those at the base, reflecting the initial DC formation (Fig 1, E7.5) and later protrusion (Fig 1 E8; SI Fig 3C-I).

What are the molecular responses following these spatiotemporally entangled physical-morphogenic events? Mechanical forces such as stretching have shown to be able to induce rapid changes in cellular behaviors such as epithelial-mesenchymal transition (EMT, a more fluidic-like behavior) by EMT-actuators such as Tgfβ, Snai1 and Slug (Leggett *et al*, 2021). Interestingly, we found that Slug, an EMT-associated transcription factor that enhances epithelial migration, is expressed at the whole placode epidermis at E7, which later shifts toward the posterior bud epidermis where the posterior bud protrusion continues on E8(Li *et al*., 2013). In the meantime, the region surrounding the DC is triggered to express molecules in ring patterns as seen in the whole mount staining (Fig. 1G-I). Sostdc1 is expressed in the epidermis surrounding the feather bud (Fig 1G). In the dermis, DKK1 is expressed at the periphery of DC. Gremlin, in a bigger ring configuration, is expressed outside of the DC (Fig 3H). By overlaying these concentric zone maps and their known properties, we interpret that Slug decreases epidermal stiffness while Sostdc1 increases epidermal stiffness, and DKK1 and Gremlin decrease dermal stiffness (Fig 1I, SI Fig 4B). This interpretation is based on previous findings that Dkk1 and Gremlin weaken dermal condensation (Chen *et al*., 2015), leading to decreased dermal stiffness. Together, after the bud protrusion induced-stretching, one anticipates a feedforward connection between these molecular signaling events and the physical-morphogenetic processes. These responses prelude the mechanical environments for epidermal invaginations that will take place in the next stage at E9.

**Fig. 3.**
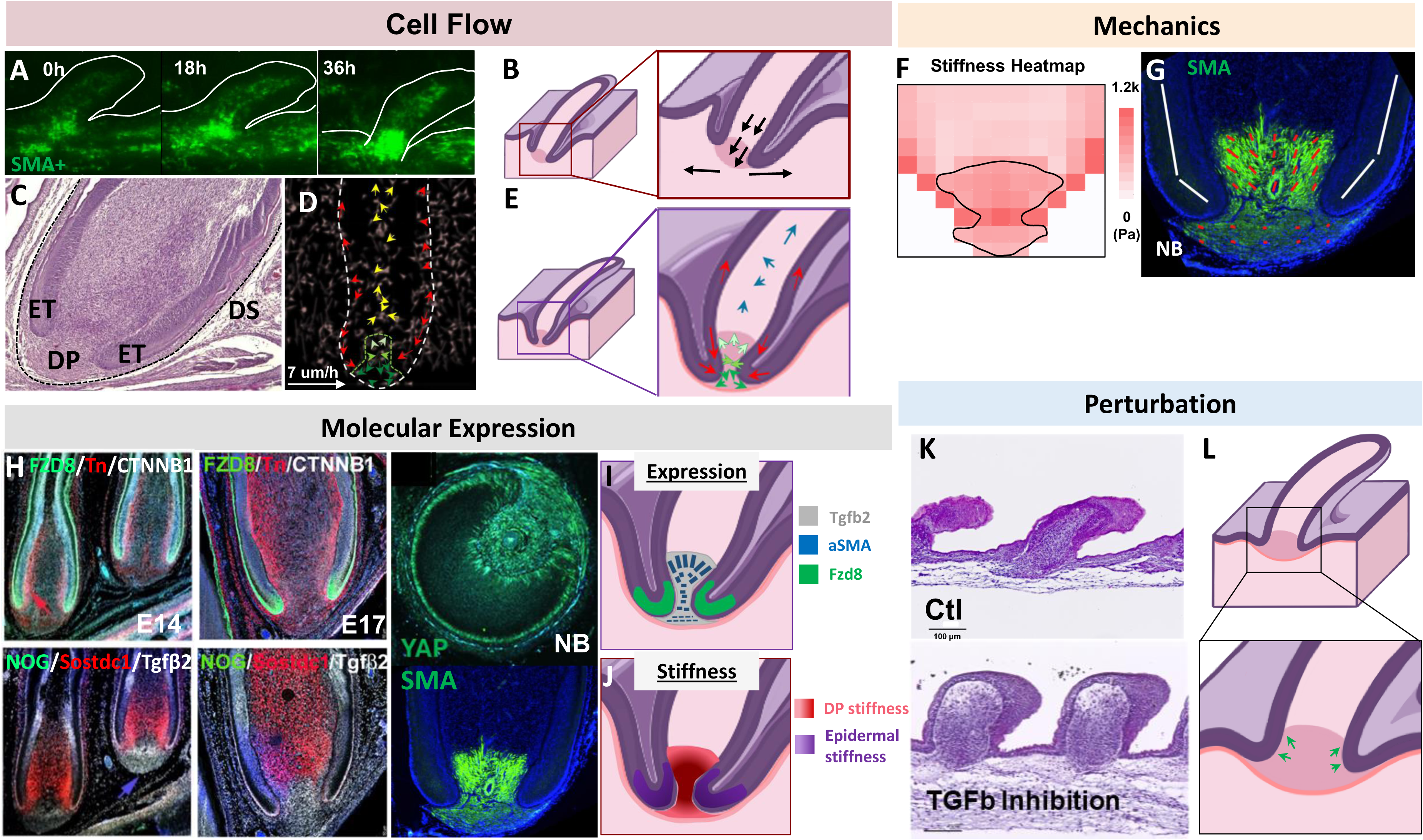
Formation of follicle base: Establishing dermal papilla through compaction of epidermal tongues on dermal condensation cells. A. Lateral view of time-lapse photos of lentivirus labelled SMA+ cells (green) in E9+36h explant. The white lines demarcate the outline of a feather bud. B. Illustration of SMA+ cell migration pattern during early DP formation. C. H&E of an E17 flight feather follicle to highlight the epidermal tongue (ET), DP and dermal sheath (DS, dotted line). D. Cell tracking of E17 flight feather to reveal differential cellular flows: (green) DP forming cells migrate downwards. Epidermal tongue cells migrate downwards while distal feather epidermal cells migrate upwards (red). Feather elongating dermal cells migrate distally (yellow). Green dotted line demarcates presumptive DP, and top (light green), middle (green) and lower (dark green) DP cells show differential migration patterns. E. Illustration of feather follicle formation and differential cell movements in late stages. Blue: feather elongation dermal cells. Red: epidermal cells. Light, green, and dark green: top, middle and lower DP cells, respectively. F. Stiffness heatmap of E17 flight feather dermal papillae. Sagittal section of follicles measured by AFM. Black lines demarcate dermal papillae. G. SMA+ cells in the DP region of a newborn (NB) feather. The orientation of SMA cells is highlighted by red lines, and expression level by length of the arrow. White line: alignment of the epidermal wall. H. Molecular expression in feather follicles. FZD8, TNC, CTNNB1, NOG, SOSTDC1, Tgfβ2, YAP (longitudinal section) and SMA expression in E14, E11, E17 and newborn flight feathers and particularly around dermal papillae region. I. Illustration of spatial expression of Tgfβ2, SMA and Fzd8 during late follicle formation. J. Illustration of DP and epidermal spatial stiffness during late follicle formation. K-L Perturbation of the follicle base formation by disruption of Tgfβ2 signaling K. H&E images of control (Ctl) and Tgfβ2-inhibitor treated explant (E12 + 4d) showing disrupted late epidermal invagination and failure of dermal condensation to compact into dermal papillae. L. Illustration of disrupted late epidermal invagination and expanded DP formation.

The observations above indicate that bud protrusion is associated with local changes in tissue mechanics. To test whether tissue mechanics can reciprocally influence the physical property of the developing feather bud, we performed various molecular functional perturbations on the mechanical properties of the epidermis and dermis. Sprty2 has been shown to enhance epithelial invagination (Schneider *et al*, 2017; Sternlicht *et al*, 2006; Yue *et al*, 2012). By overexpressing Sprty2, we observed a decrease in epidermal stiffness (from 1.62 ± 0.18 to 0.91 ± 0.10 kPa) (Fig 1K), associated with a visibly widened distal end and a shorter bud length in feather buds (Fig 1J). These results suggest that epidermal stiffness helps confine the bud shape, and the softened epidermis may allow the dermal cells to migrate laterally, leading to the expansion of bud width (Fig 1L). Mechanistically, cell flow and tissue mechanics both depend on cell contractility and adhesions. We treated the explants with Blebbistatin – a known inhibitor of non-muscle myosin II and therefore cell contraction and migration (Wu *et al*, 2019). This led to a sac-like bud morphology. This may be due to the failure of dermal cells to migrate vertically upward and create reciprocal forces at the epidermis (Fig 1M, SI Fig 4C). Similarly, treating cells with ADH-1, a peptide that blocks N-cadherin (Shintani *et al*, 2008), which is enriched in the DC, led to the formation of shorter feather buds (Fig 1N)., suggesting that dermal cell-cell adhesion is an important factor contributing to the forces driving vertical movement during short bud formation. Together, these results show feather bud protrusion is achieved through a sequence of spatial biomechanical-biochemical interactions and characterized by dermal cells to flow vertically and upward, while keeping the interbud skin in place.

#### Bud elongation stretch-activates epidermal YAP and MMP in a ring around the bud-base and initializes downward epidermal folding to form the stiff follicle wall

During the follicle formation stage (E9-12), the epidermal cells surrounding the bud base invaginate downward into the dermis (Fig. 2A, B, red arrows), folding into a bi-layered cylindrical wall around the presumptive dermal papilla (DP). Cell tracking analysis shows that during epidermal invagination, while the feather bud continues to elongate at the distal end to form putative pulp (Fig 2A, B, yellow arrows, Supplementary Video 2), dermal cells at the bud base turn their migration direction downwards and form the putative DP (Fig 2B, green arrows). These dermal cells condense as the distance between neighbouring cells decreases, and express SMA (α-smooth muscle actin) (SI Fig 5A-B), implying higher cell compaction and stiffness (Late invagination. green arrows, SI Fig 5C, cyan line). Outside the invaginating wall, dermal cells flow along the epithelial fold, forming the putative dermal sheath (DS) (Fig 2B, blue arrows). The illustration in Fig 2C shows how the epidermal invagination (red arrows) forms a cylindrical wall around the putative DP, and at the same time generates new localized cell flows along different parts of the follicle wall (Fig 2C, lower panel). Thus, we interpret that the mechanically rigid invaginating follicle wall (Fig 2D) serves as a physical-mechanical barrier to compartmentalize and organize the mechanical signaling of dermal cells as well as epidermal cells, and to specify new cell fates within and outside of the feather follicle. From a mechanical perspective, it is also likely that migrating cells with a certain degree of stiffness tend to invade softer regions rather than stiffer ones.

In line with this interpretation, Martino et al. have shown that dermal sheath-generated contractile forces orchestrate homeostatic tissue regression (Martino *et al*, 2023), which also implies that SMA+ cells – abundantly observed during follicle development - could be an important source of force generator, participating in creating the differences in tissue stiffness. Additionally, the stiffness of the bi-layered invaginating epidermal cells is more than twice stiffer (1.04 ± 0.11 kPa) than the surrounding dermis (0.42 ± 0.03 kPa) (Fig 2D), providing the mechanical feasibility for the epidermal cells to invaginate into the softer dermis around the bud. The moderately stiff DC (0.72 ± 0.07 kPa) could also serve as the physical barrier that divides distal bud dermal cells (0.32 ± 0.03 kPa, to become pulp) from the basal dermal cells (0.44 ± 0.05 kPa, to become DP). This mechanical contrast suggests the role of high stiffness cells (invaginating epidermal cells, DC cells) serving as physical barriers to compartmentalize cell fates during feather follicle formation.

One consequence of cell migration is the generation of mechanical influences on the surrounding cells. To assess the mechanical cues associated with the initial localized chemo-mechanical remodeling events that lead to epidermal invagination, we performed QMorF analysis to survey the forces exerted around the bud base based on the regional cellular deformation (Fig 2E-G). We observed that as the feather bud continues to elongate, it stretches the epidermal cells at the neck of the feather bud adjacent to the DC (Fig 2F, white dotted circle), which could activate other chemo-mechanical coupling molecules. More specifically, epidermal invagination is characterized by high-order 3D morphological transformations (SI Fig 3J-M), which can be appreciated by an azimuthal approach (Campas & Mahadevan, 2009), in which we treat the 3D morphological deformation of the cells in a concentric perspective by using the center of the feather bud as the central axis. Analysis using our Azimuthal (using the dermal condensate as the center point with respect to epidermal invagination) approach indicates that the mechanical coupling and the associated cellular deformation accommodate the layering of cellular elements into the azimuthal symmetric bud in the 3D configuration.

To form an azimuthal invaginating neck around the bud, a cellular band composed of small elements to azimuthally shrink the physical dimension around the original bud cylinder near the invagination site is a straightforward approach. This argument is supported by the orange band in Fig. 2F, representing a region surrounding a bud with a relatively smaller cellular area than the rest between two gray dashed curves in Fig 2F. The opposing arrows on either side of the small-cell band can be considered as a force field interface of two contributing to the distinctive 3D invagination of a follicle macroscopically (Fig. 2F-G and SI Fig 3). In summary, our QMorF results demonstrate that the morphogenesis of the 3D follicle is achieved through a series of morphing processes across scales and response to mechanical signaling organizers - molecules that respond to and cause mechanical changes leading to morphogenesis.

We then search for molecules related to mechano-chemical signaling and involved in epidermal invagination. Stretch-activated YAP has been shown to mediate such chemo-mechanical coupling (Halder *et al*, 2012). Here we found YAP is highly induced in the bud base and especially in the invaginating epithelial walls (Fig. 2H), where MMP activities are also elevated (Fig 2H) in contrast to the apteric region (SI Fig 5D). Gremlin and Sostdc1 were also observed at the invaginating dermis and epidermis, respectively (Fig 2H). These results could imply that, initiated by feather bud elongation induced-stretching, YAP, Wnt, MMP, and others work in concert to soften the dermis around the DC and facilitate epidermal tongue invaginations (Fig 2D, I).

To test the role of follicle wall stiffness in regulating DC morphology, we overexpressed Snail and observed lowered follicle wall stiffness, widened buds with a broader base, and failed follicle invagination (Fig 2J). Wnt3a overexpression resulted in failed invagination characterized by shortened and widened feather bud formation. Furthermore, longitudinal-sectional view of Wnt3a overexpressing feather follicle shows that without epidermal invagination, the physical constraint for DC cells is lost, leading to “dispersed” dermal cell flows (Fig 2K). MMP inhibition suppresses ECM remodeling and has been shown to suppress follicle invagination(Jiang *et al*, 2011). Together these results suggest that follicle formation is achieved via stretching of the bud / interbud boundary – indicated by a stiff epidermal wall, YAP expression and a stretched cell morphology - that triggers the initial epithelial invagination. The process is characterized by adequate mechanical control that keeps the epithelial tongue extending downwards continuously, which in concert with dermal cell migration, transforms the 2D planar skin into a 3D vertical follicular architecture (Fig 2L).

#### The stiff epidermal wall invaginates and compacts the Tgf-β emitting dermal papilla to form the base of the follicle and dermal papillae

To complete follicle formation (E17 in flight feather), we observe the epidermal wall (white line, Fig 3A) reaches down to the follicle base to clamp around the DP (SMA+ cells, Fig 3A), while being wrapped around by the DS from the outside (Fig 3A-C). We analyze the differential cellular flows and tissue rigidity changes in this final stage. Cell tracking analysis reveals new complex cellular flows that emerge at this stage (Fig 3D, Supplementary Video 3). Epidermal wall cells at the base (lower red arrows) migrate downwards and bend towards the putative DP, while epidermal cells above the putative DP migrate upwards (upper red arrows), along with the dermal cells within the follicle (yellow arrows). The migration of putative DP cells (green arrows) is relatively slow. This flow can be further divided into 3 sub-flows: below the epidermal wall (dark green arrows, Fig 3D-E), on par with epidermal wall endings (green arrows, Fig 3D-E), and inside the epidermal wall (light green arrows, Fig 3D-E). DP cells below the epidermal wall mostly migrated horizontally and slightly downwards, those on par with the epidermal tongues migrating inwards horizontally, while those inside the follicle wall showed moderate upward movements (Fig 3D-E, green arrows). The illustration shows the intricate movement of cell flows during DP formation (Fig 3D), which are parallel to the cell movement during feather cycling in the adult feathers (Wu *et al*., 2021b).

The landscape of tissue stiffness of the follicle is surveyed using longitudinal sections of the follicle and AFM. The epidermal wall and invaginating tongues show the highest stiffness (1.22 ± 0.13 kPa), followed by DP. Specifically, the bottom section of DP where the epidermal “tongues” meet the DP shows the highest regional stiffness, suggesting the epidermal structures not only act as physical barriers but also as physical “stressors” that compacted and seal the follicle floor with DP (Fig 3E). High regional stiffness coincides with cell density, which includes DP (SI Fig 5E). SMA is highly expressed by the DP cells in the upper DP (Fig 3F) and is in alignment with the pattern of observed cell flow (Fig 3D), radially dispersing from the narrowest part of the DP. Beneath this follicle floor, dermal cells express moderate levels of SMA and are circumferentially and horizontally oriented (Fig 3F), forming the DS cup to further seal the follicle floor.

These results imply that differential dermal cell flows, through interaction with epithelial tongues, are compartmentalized to form DS along the outside follicle wall, pulp cells along the internal follicle wall, and DP and DS cup at the bottom. Pulp cells and DS cells are oriented longitudinally, while DP and DS cup cells are oriented circumferentially. Thus, cell flows are locally distinct and correlate with the difference in tissue stiffness. Dermal cell orientations and cell flows in upper and lower regions of DP suggest mechanical factors play an important role in dermal cell behavior and fate specification (Fig. 3G, H).

We then ask what the molecules that mediate these biomechanical – biochemical interactions are to complete epithelial tongue invagination and DP formation. Tgfβ2 is known to induce mesenchymal condensation (Ting-Berreth & Chuong, 1996) and can also activate SMA, induce cell contraction and EMT(Massague & Sheppard, 2023). Here we found that Tgfβ2 is expressed by the dermal cells at the base of the follicle starting at E14 and through E17 (Fig 3H); this corresponds adequately to the high expression of SMA in the DP and Fzd8 at the tip of epidermis. In the prostate gland, Tgfβ-induced signaling is regulated in part by Wnt receptor Fzd8 (Spanjer *et al*, 2016). These results suggest that DP may use Tgfβ2 as a chemotaxis cue to attract epidermal invaginations, indicated by Fzd8 expression. The high expression of YAP at the tip of the epidermal tongue and DP in NB flight feather DP region also suggests the DP cells and epidermal-DP interface are experiencing high level of mechanical stress – potentially shaping DP into its final form (Fig 3G).

To test whether Tgfβ2 indeed functions as a chemoattractant to the invaginating epidermis and also shaping of DP, we suppressed its activity by treating the explant with LY2109761(Melisi *et al*, 2008), a Tgfβ I and II kinase inhibitor, from E12 to E16. The suppression of Tgfβ2 activity reduced epidermal invagination, and the narrowing of DP at the follicle base was lost (Fig 3H). The results imply that by inhibiting Tgfβ2 signaling from E12-16, the chemoattracting signal from the DP to the invaginating tongues is stopped, and hence the “DP shaping” role of the epidermis could not be completed, leading to a loosened DP and widened follicle base (Fig 3I). Inhibition of Tgfβ2 seemed to also hamper feather elongation (shortened feather bud), which could be an important mechanical cue that initiates epidermal invagination (Fig 2).

These results suggest the process to close the base of feather follicle is characterized by a series of mechanically and chemically interconnected events to achieve mechanically guided cellular flows, tissue folding, and finally cell fate specification. Molecular and tissue mechanics work in concert to induce shape changes. When putative DPs reach a compaction threshold, they secrete more Tgfβ that trigger the bending of the epithelial tongue toward DP, ending the follicle elongation process and closes the bottom of the follicle with DP and DS cup. In so doing, other molecules are induced in different topological positions in this newly set up proximal follicle, positioning themselves for future cyclic renewal (Wu *et al*., 2021b). These include Tenascin C (TNC) in the dermal cells adjacent to invaginating epidermal walls, as well as NCAM and some extra-cellular matrix (ECM) molecules (Chuong & Edelman, 1985), and Sostdc1 expressing dermal cells which are now inside the feather follicle and distal to DPs (Fig 3G).

#### From 2D plane to 3D follicle: A mechano-chemical coupling model for topological transformation during follicle formation

Upon dermal condensation formation, we observed a switch in dermal cell movement associated with changes in epidermal cell shape and stiffness. These events were followed by epidermal invagination around the DC, which ceased when the invagination reached the base of the DC (Fig 4A). We wonder whether the causality of these spatiotemporally entangled events can be addressed by a simple mechano-physical principle. Mechanically, condensed dermal cells at the DC exert traction forces on the surrounding tissue. While the forces propagating horizontally or vertically downwards can be counteracted by the traction forces from the surrounding DCs or the resistance from the dermis, the forces propagating vertically upwards would eventually reach the epidermal layer and act as downward pulling forces on the epidermal cells. Meanwhile, dermal cell-secreted chemical factors, such as Tgfβ, can stimulate epidermal cells and induce their morphogenetic movements. Tgfβ can also signal the surrounding dermal cells to increase the secretion of MMP (Moore-Smith *et al*, 2017), which further leads to the activation of Tgfβ (Kobayashi *et al*, 2014). As MMP can degrade the basement membrane (BM) (Strzyz, 2019), the synergistic effect of Tgfβ and MMP promotes the formation of mechanically weakened regions in the BM near the DC region to facilitate the invagination of epithelial cells. We wonder if these dermal mechano-chemical activities can address the changes in epidermal cell shape and stiffness at the early stage of bud protrusion, particularly the weak point formation that allows the switch of dermal motion, the specification of invagination site, and the termination of invagination. Investigating these possibilities is not trivial if we use an experimental approach. As such, we instead used coarse-grained phenomenological modeling approach (Fig. 4B).

**Fig 4.**
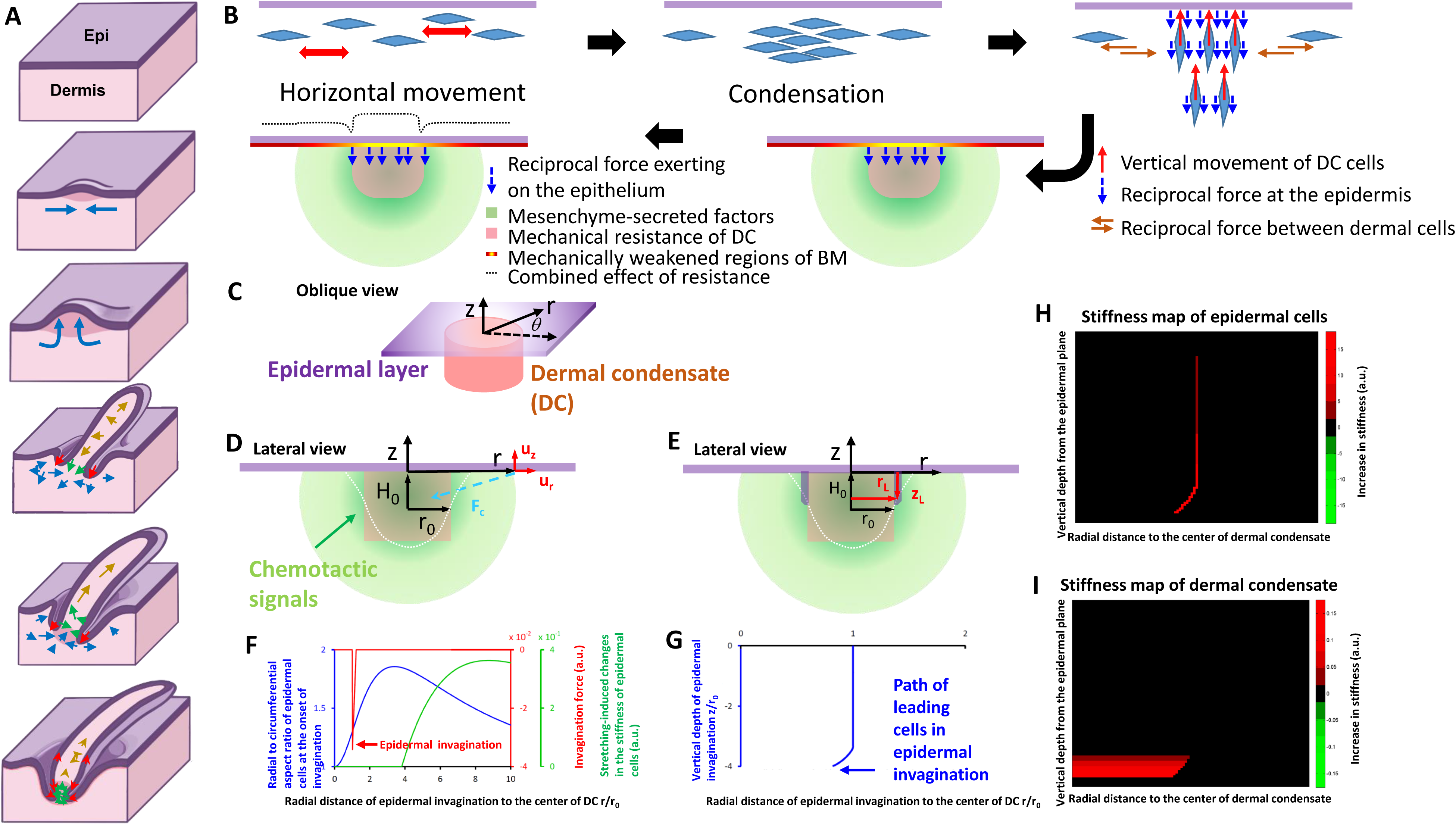
A mechano-chemical coupling model of feather follicle formation. A. Morphological transitions of the topological transformation are shown in five steps. a, b are studied in Fig. 1, c, d are studied in Fig. 2. e is studied in Fig. 3. Through these inter-connected processes, a complex follicle is formed from the planar epithelium. B. The schematic depicts the model assumptions. Following the horizontal movement, the FGF-stimulated dermal cells form dermal condensates (DCs) and generate migratory/contractile forces therein. While the forces in the horizontal (orange) or downward (red) directions are counteracted by the forces from neighboring condensates or the resistance from the deep dermis, the forces in the upward (red) direction create reciprocal traction forces at the epidermis (blue). Meanwhile, the diffusive Tgfβ-MMP signaling (green) creates a weakening effect (yellow) at the basement membrane (BM), where the epidermal cells experience the largest mechanical resistance (light pink) from the DC. The model aims to investigate whether the combination of BM weakening and DC resistance (dotted gray line) and the diffusive Tgfβ-MMP signaling (green) can: a) induce changes in cell shape and stiffness as observed in the experiment, and b) facilitate epidermal invagination at the periphery of the DC. C. The model simplified the epithelial layers as a flat, planar sheet (Purple) and the dermal condensate as a cylindrical structure (Pink) with a radius denoted as “*r*_0_.” The chemo-attractants (Green) secreted from the dermal cells to induce epidermal invagination were approximated as originating from a point source at a distance denoted as “*H*_0_” from the epidermal layers. A cylindrical coordinate system (*r*, *θ*, *z*) with azimuthal symmetry was employed to model the local deformation of cells within the epidermal layers, represented as (*u_r_*, 0, *u_z_*), during the onset of chemoattractant-induced invagination. **F-G**) The numerical results for panel **C**) indicate an increased radial to azimuthal aspect ratio (Blue) and enhanced cell stiffness (Red in panel **F**) resulting from cell stretching in epidermal cells located distantly from the dermal condensate. The maximal force for invagination was observed near the border of the dermal condensate (Red arrow in panel **B**). **E**) The model used the trajectory of epidermal leading cells at the forefront of invagination, denoted as (*r_L_*, *θ*, *z_L_*) with *θ* ranging from 0 to 2*π*, to depict the progression of invagination along the sidewall of the dermal condensate and the subsequent closure of invagination at the base of the newly formed follicle. **H-I**) The numerical results show **G**) the path of the invagination (Blue) and the increase of stiffness represented as heat maps for **H**) the epidermal cells and **I**) the dermal cells during invagination. The parameters utilized in the numerical simulations are detailed in the model section.

At the epidermal-dermal interface, we assumed that the epidermal layer received both a traction force and a chemo-attractive force from the underlying DC (Fig. 4B, the blue lines and the green marks). These epidermal cells will encounter resistance from the DC if they start moving downward. As a result, only epithelial cells around the peripheral border of the DC move downward, thereby specifying the ring-like zone of invagination. The region with the highest Tgfβ-MMP signaling activity is presumably located near the base of the DC, prompting the epidermal cells to move towards this target through invagination. Initially, there is a considerable distance between the target and the epidermal cells. The force generated by the Tgfβ-MMP signaling is insufficient for the epidermal cells to deform the DC, causing the epidermal cells to move along the periphery of the DC. As the epidermal cells move towards the region with the highest concentrations of chemo-attractants, the force increases, allowing the cells to compress the DC, resulting in the closure of the invagination. In terms of changes in stiffness, the epidermal cells and dermal cells rely on distinct mechanisms. Dermal cells experience an increase in stiffness primarily due to the compression of ECM molecules, a phenomenon previously demonstrated to enhance stiffness (Kim *et al*, 2017; Tronci *et al*, 2013; Vinciguerra *et al*, 2014; Wollensak *et al*, 2003). This compression results from the centripetal migration and/or constriction force of the epidermal cells along the DC boundary, which also causes the crowding/narrowing of the DC and the convergent alignment of dermal cells near the base of the DC. On the other hand, epidermal cells change in stiffness through two distinct processes. When located away from the target where Tgfβ-MMP signaling activity is low, the stiffness of epidermal cells primarily increases due to elastic cell stretching within the epidermal layers. Conversely, when epidermal cells are near the base of DC, where Tgfβ-MMP signaling activity is high, cell stiffness increases even in the absence of chemoattractant gradients (Gladilin *et al*, 2019).

Using these assumptions, we constructed a simple model to address the biophysics of the morphogenetic events associated with the onset and closure of epidermal invagination. The model focuses on the invagination and neglects the protrusion of the bud. For the onset of invagination, we focused on the elastic responses of the epidermal cells to the dermal traction and chemo-attractive forces. Using a cylindrical coordinate system (Fig. 4C-E), we analyzed the changes in epidermal cell aspect ratio (Fig. 4F, blue curves), the position to initiate invagination (Fig. 4F, red curve), and the changes in epidermal cell stiffness (Fig. 4F, green curve) through a steady-state approach. We observed that the aspect ratio increases gradually with the radial distance *r* from the center of the DC, and then gradually returns to the baseline upon reaching the inter-bud region. In comparison, cell stiffness increases in the region away from the DC due to significant cell stretching in that area. For the closure of invagination, we focused on the movement of leading cells located at the forefront of the invagination (Fig. 4E). A centripetal moving pattern of the leading cells was observed near the base of the DC (Fig. 4G). Accordingly, the stiffness of these leading cells gradually increases as they approach the base (Fig. 4H), with a simultaneous local increase in the stiffness of the DC due to the centripetal closure of epidermal invagination (Fig. 4I). As such, the model adequately describes the force distribution and deformation of epidermal cells caused by mechano-chemical cues from the DC to initiate, extend, and complete epidermal invagination and DP formation. More details can be found in the Methods section and the Supplementary Information (SI).

#### Scale to feather conversion: softening scale epidermis enables feather follicle formation

Among the many amniote skin appendages (Wu *et al*., 2004), only few form follicle configurations that allow cyclic renewal, including hairs (Fuchs, 2009), feathers and alligator teeth (Wu *et al*, 2013). Scales represent epithelial folding and contain epidermal progenitors that allow scales to grow, but not molting and regeneration. The unique aspect on birds is that scutate scales can be converted into feathers (Dhouailly, 2009; Widelitz *et al*, 2000; Wu *et al*, 2018a; Wu *et al*., 2018b). Here, we revisited the scale-to-feather transition (Lai *et al*., 2018; Wu *et al*., 2018b) with a new bio-mechanical perspective on their cell flows and tissue rigidity. Scutate scale is characterized by a flat protrusion off the skin surface and with limited posterior elongation (Fig 5A). Cell tracking analysis during scutate scale development shows that the epidermal cells travel from anterior to posterior, while dermal cells show limited posterior and vertical migration (Fig 5B, Supplementary Video 4). This suggests the relatively “soft region” of the scale could be at its posterior end. The stiffness of the scutate scale is 23% higher (1.63 ± 0.17 kPa) at the anterior than the posterior protruding edge (1.23 ± 0.11 kPa, Fig 5D), and both are stiffer than that of a feather bud (0.84 ± 0.09 kPa, Fig 2A). The generally rigid scale also implies the dermal cell flow is limited and little “stretching-induced” mechanical signaling could occur.

**Fig. 5.**
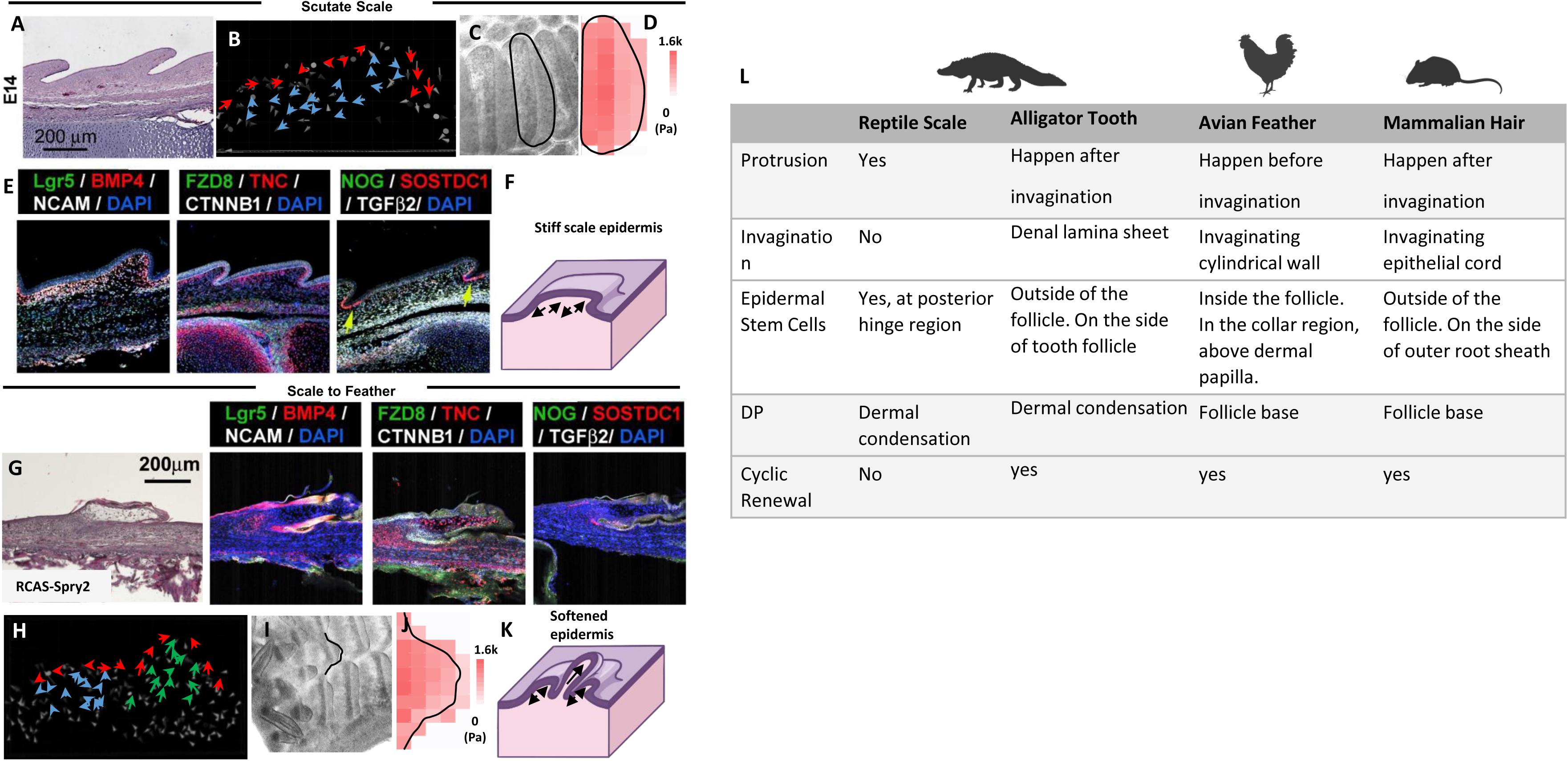
Scale – feather transition: Biophysical events following biochemical signaling alterations re-direct morphogenetic consequences. **A-F: Biophysical characterization of scutate scales and their formation.** A. H&E of E14 scutate scale B. Cell tracking analysis of E10+24h quail scale showing limited dermal cell migration during scale formation C. Bright field top-down view of scutate scale in E14 explant. The black line demarcates the scale measured by the AFM D. Stiffness heatmap of an E14 scutate scale. E. Lgr5, BMP4, NCAM, FZD8, TNC, CTNNB1, NOG, SOSTDC1, and Tgfβ2 expression in E14 scale. F. Illustration of stiff scale epidermis limiting dermal cell flow. G-K: Scale-feather conversion is accompanied by epidermal softening and redirected dermal cellular flows. **G. H&E section and molecular expressions (shown by RNAscope) of scale to feather conversion by overexpressing Spry2 in the epidermis.** H. Cell flow analysis of newly induced feather buds on top of the scutate scale. It shows dermal cells are re-directed to generate vertical flow in the newly formed feather bud. I. Bright field, top-down view showing scutate scale-to-feather conversion in spry2 overexpressing explant. The black line demarcates the scale measured by the AFM J. Stiffness heatmap of an E14 scutate scale-converted feather buds. K. Illustration of scale to feather transition showing softened epidermis on the scutate scale allows dermal cells to flow vertically and generate feather buds. L. Comparison of cellular flows during the formation of reptile scale, avian feather buds and mammalian hair primordia. Bud protrusion, follicle invagination and dermal papillae formation are compared.

We wonder which biochemical circuit can transduce biomechanical force here. While Tgfβ2 is expressed in the basal layer of the scale dermis (Fig 5E), FZD8 is not significantly expressed in the scale (Fig 5E). NCAM is expressed mainly at the anterior end of the scutate scale, TNC is expressed is expressed in the dermal cells located at the epidermal-dermal interface (Bao *et al*, 2016), and SOSTDC1 is only expressed at the epidermal “anchors” of the scale, in contrast to its high expression in the DP cells during feather follicle development. The high expression of dermal cell adhesion molecule NCAM and TNC at the dermal-epidermal interface suggests strong dermal adhesion and epidermal-dermal binding. High epidermal stiffness at the anterior end probably helps to guide the limited dermal cell migration posteriorly, where the epidermal cells are slightly softer and enable some protrusion to occur (Fig 5F). Yet, the lack of a signaling center that mediates biochemical and biomechanical cross talks in the scale prevents changes in spatial tissue stiffness, dermal cell mobility and epidermal invagination into the dermis.

Can we alter the tissue mechanics of the scale to allow feather follicles to form on the scale? Can softening the scale epidermis by Spry2 facilitate this transition? Our previous study(Wu *et al*., 2018b) shows that locally overexpressing Spry2 in the epidermis can convert scale to feather (Fig 5G). Cell migration analysis shows that during Spry2 overexpression-induced scale to feather conversion, the vertical dermal movement observed in feather bud formation is also observed at the converted feather bud (green arrows, Fig 5H, Supplementary Video 5), accompanied by 65.6% softening of the local epidermis (from 1.62 k to 0.48 kPa, Fig 5I, J). We interpret the result as epidermal Spry2 overexpression softened the epidermis and allowed dermal migration to overcome the physical barrier of the epidermis to form a feather bud-like structure (Fig 5K). In comparison, the reptile scales are characterized by short and wide dermal protrusion, limited epidermal invagination, and no dermal papilla or follicle-like structures (Di-Poi & Milinkovitch, 2016). The mammalian hairs do form follicles and exhibit robust cyclic renewal behavior (Fuchs *et al*, 2001; Stenn & Paus, 2001). In development, hair germs invaginate deep into the dermis, and the follicle wall wrap around DP through different morphogenetic processes (Morita & Fujiwara, 2022). Eventually, hair stem cells are located in the bulge region (Li & Tumbar, 2021; Morrison & Spradling, 2008), different from the collar bulge in feathers(Yu *et al*., 2002). While both hairs and feathers cycle effectively, the different topology and formative processes imply convergent evolution of their follicle formation processes (Fig 5L).

### Discussion

We propose that a morphogenesis transition circuit requires a sensor and actuator to undergo phenotypic changes (Fig 6A), and the chemo-mechanical coupling events of a developing system are a demonstration of sensor – actuator self-organizing signaling circuit (Fig 6B). A morphogenetic system senses and responds via a sensor and actuator mechanism to achieve its respective morphogenesis stage. Once actuated, it becomes the sensor for the following stage. During feather follicle development, an originally homogenous skin tissue undergoes 1) bud protrusion, 2) invagination to reach 3) follicle formation (Fig 6B). Any incompletion of the previous stage would not lead to the initiation of the following one.

**Fig. 6.**
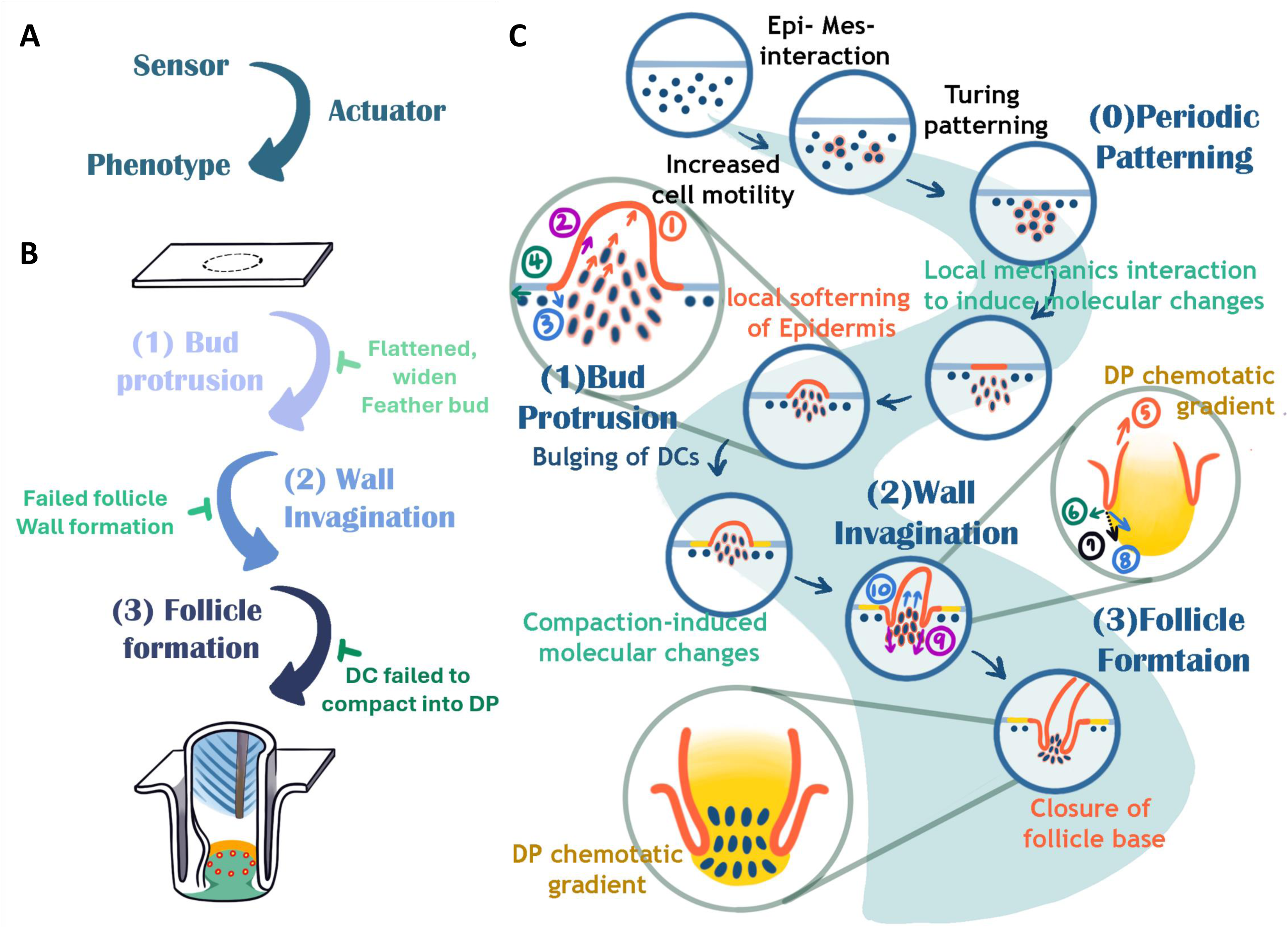
Schematic illustration showing a series of self-organizing cellular flows, driven by mechano-chemical coupling, can build the complex topology of feather follicles. A. self-organizing signaling circuit. The circuit requires a sensor and actuator to undergo phenotypic changes. This is the core circuit. B. Multiple self-organizing signaling circuits can be coupled to achieve a more complex morphogenetic process. The coupling happens when the product of one process become the initiator of the next process. C. Here, the formation of feather follicle is used to illustrate the coupling process. During feather follicle development, an originally homogenous skin tissue undergoes Turing periodic patterning to generate a population of feather primordia. Here we focus on one primordium. We study how an individual 2D placode is transformed into a 3D follicle architecture. We highlight three stages of morphological transition: 1) bud protrusion, 2) invagination to reach 3) follicle floor formation. The involved mechanical forces, some are hypothetical, are shown in arrows: (1) Protruding stress, (2) Cell flow-induced shear stress, (3) Chemotactic force, (4) Inter-bud skin tension, (5) Epidermal proliferation dermal cell flow-induced epidermal migration, (6) Elastic resistance force from the DC, (7) Net force, (8) Chemotactic force, (9) Expansion force from cell sheet + ECM-induced migration force, (10) Rebounding force from compacted dermal cells. Epi: epidermis. Mes: Mesenchyme. DC(s): dermal cell(s). WP: weak point.

The result is summarized in Fig. 6C. The topological transformation during feather follicle formation is best described in patterning, protrusion, invagination, and follicle floor formation stages. The three stages are coupled in a way that the consequence of one stage becomes the initiator of the next stage. This is evident in the case of scale to feather conversion, where softening the epidermis not only leads to the initial feather bud formation, but also the sequential epidermal invagination and eventual follicle architecture (Fig 5).

*Stage 1, Patterning*: <u>Cause:</u> dermal condensation process leads to increased stiffness in its center. The dermal cells need be pushed either downward into the dermis (as seen in hair follicle formation) or protruding up (as seen in the feather). In the feather case, the placode epidermis becomes softer (as evidenced by AFM measurement). <u>Results:</u> Vertical dermal cell movement upward, making bud protruding out. Please also see point 2 for the dermal condensation process.

*Stage 2, Invagination*: <u>Cause:</u> As the bud grows distally, the epithelial cells at the bud base (in a ring configuration) is stretched (judged by cell shape change and expression of YAP, Sostd, MMP). <u>Result:</u> The epithelia (in a cylinder configuration) surrounding the bud invaginate into the dermis

*Stage 3, follicle floor formation*: <u>Cause</u>: As the invaginating cylindrical follicle wall and the dermal condensates within continue to dig into the dermis (Fig. 3A-E), we observe dermal condensation cells start to express SMA, exhibit circular cell arrangement (Fig. 3G, H), become stiffer (Fig. 3F), “pulling” adjacent fibroblast and make them express YAP (Fig. 3H), and expressing a higher amount of Tgf-β (Fig. 3H) that attracts the invagination epithelial wall to bent toward the dermal papilla. <u>Results:</u> the formation of dermal papilla and sealing of the follicle base.

Our previous work has shown that optimal tissue stiffness is required for regenerative wound healing to occur(Harn *et al*, 2019; Harn *et al*, 2021), in which a soft dermal environment allows stiff epidermal hair placodes to invaginate downwards and form the hair germs. Proper epidermal stiffness is also required for the development of hair follicles (Jamora *et al*, 2003). Similarly, this also suggests that there should be an optimal stiffness for the epidermis and the dermis for each stage of feather development. Specifically, the stiffness contrast – a force imbalance between the local epidermis and dermis leads to tissue folding, guides cell flow and hence morphogenesis. At E7, the actively dividing and migrating dermal cells create enough stochastic events to increase random cell movement – enhancing cell-cell adhesion and the emergence of the initial DC cells (Fig. 6). This process increases DC stiffness from ∼175 Pa to ∼212 Pa. These local mechanical changes in the DC can interact locally with the epidermis, demonstrated by the aspect ratio and orientation of the epidermal cells, local Shh/Slug activation and softening (<196 Pa), and allow dermal cells to bulge through, forming the protruding short feather bud. Snail, a key molecule identified in softening epidermis, is a known downstream molecule of Shh signaling pathway (Riaz, Ke et al. 2019), which explains the often colocalizing expression pattern of these 2 molecules (Fig 3A). Snail is known for downregulating epidermal cell junctions (Cano, Perez-Moreno et al. 2000); hence, it is reasonable to observe the overall reduction in epidermal stiffness when it is expressed.

On the dermal side, the FGF promoted “solidification” of DC core has been proposed to work with contracting surroundings enabled by BMP to drive the budding process (Yang *et al*., 2023). We should take this opportunity to clarify that here we are focusing on the protrusion from DC to short feather buds (bud base larger than height), while the elongation from short to long bud is controlled by Shh/ Gap junction dependent calcium activity and dermal cell migration(Li *et al*, 2018).

As bud continue to grow distally, the dermal cell flow not only facilitates feather bud elongation, but also causes shear stress and stretches the epidermis surrounding the bud base (Fig. 6). This in turn activates epidermal YAP and its downstream signaling events: proliferation, cell motility, and MMP, leading to the stiffening of epidermis (∼1 kPa), and enabling them to “fold and invaginate” through a buckling process into the softened local dermis (∼0.5 kPa), followed by a chemoattractant-mediated extension of the invagination. YAP is a well-known mechanically-activated molecule(Panciera *et al*, 2017), and its activity at the epidermal invaginating tongues suggests the stiffening of the local epidermis, which is essential for remodeling the dermis and facilitates invagination (Fig 3E). YAP also activate MMP(Lei *et al*, 2023), which further soften the invagination region (Jiang *et al*., 2011). These chemo-mechanical molecules controlling epidermal-dermal mechanics form multiple concentric ring patterns (Fig. 1I) that resemble the telescope model proposed by Fujiwara (Morita *et al*, 2021). We postulate that these “ring-like” molecular expression play an important role in creating the “mechanical contrast” that initiates and guides fluidic cells to flow into the dermis, as observed during invagination.

In addition to feather follicle development, the scutate scale study demonstrates the importance of tissue folding and compartmentalized cell flows for cell fate specification. The limited dermal cell flow pattern implies that the morphogenesis of scutate scale is confined within one rigid epidermal compartment (Fig 5B). On the contrary, overexpressing Spry2(Wu *et al*., 2018b) softened the epidermis, which allowed dermal cell to flow, protrude and activate sequential circuits to form an ectopic feather follicle (Fig 5G, L). The ability to form DP in both avian feather and mammalian hair by extending epidermal invaginations deep into the dermis to wrap around a group of specified dermal cells, is a testament of successful convergent evolution to form stem cell-based follicles that allow cyclic renewal, enabling functional adaptation to diverse eco-spaces.

Based on this study, we propose a novel “mechanical signaling organizer” concept which works operates in parallel with “molecular signaling organizers.” A molecular signaling center typically refers to the diffusion of morphogens from a specific location, where the resulting gradient influences cell movement or cell fate. Here, we hypothesize that mechanical forces can also exert signaling effects. A mechanical signaling center would be a stiff or soft region from which mechanical forces can be transmitted, either through changes in force or through channel activities, reaching adjacent tissues more quickly and over longer distances. Recognizing the presence of these mechanical signaling organizers allows us to better understand the crosstalk between biophysical and biochemical signals. In this study, dermal condensation, the growing tip of the feather bud, and the compacted dermal papilla function as mechanical signaling organizers. These structures initiate or redirect cellular flows, enabling the formation of complex organ architectures.

In summary, this work reveals that there are unidentified key roles of dynamic cellular flows under the skin waiting to be discovered in development, regeneration and organoid morphogenesis (Lei *et al*., 2023). For each morphogenetic process, we contemplate a sensor and executer module is required to initiate and end a process. Diverse amniote scale configurations have evolved, but they all lack the follicle architecture that will allow them to undergo stem cell based cyclic renewal. Here, we demonstrate that a simple change in tissue mechanics can lead to the emergence of novel “mechanical signaling organizers,” which work alongside “molecular signaling organizers” to redirect cellular flows. When optimal conditions are met, the outcome of one morphogenetic process will trigger the next, and the sequential coupling of morphogenetic processes is initiated, as illustrated by the three consecutive stages of feather follicle formation (Fig. 6A-C). Dating back to feathered dinosaurs and Mesozoic birds, the newly evolved follicle design allowed the functionality of diverse feather forms to be tested in different skin regions and across various life stages, enhancing the adaptability of the integumentary interface to external environmental pressures. We further propose that newly generated “mechanical signaling organizers,” such as those formed during physiological molting or wounding in adults, can also redirect cellular flows to achieve repair and/or regenerative outcomes to rebuild the integumentary organ.

### Materials and Methods

**Avian skin explant model** was performed according to Jiang et al(Jiang *et al*, 2023).

**Atomic force microscopy** was used to measure tissue stiffness. Data analysis and heatmap generation were done in reference to Harn et al(Harn *et al*., 2021). In brief, a cylindrical tip with 5 um end radius and 0.2 N/m spring constant was used. Force was set at 5 nN.

#### Live imaging and cell Tracking Analysis

Live xyzt imaging was done with a confocal microscope (Stellaris 5, Leica) taking 50 um thick z-stacks of 10 min intervals for up to 24h using a 10x air lens. The sample was incubated at 37 °C, 5% CO2 and 90% humidity chamber (Oko-Lab). Imaris (Oxford Instruments) was used for cell migration analysis.

#### QMorF analysis

E-cadherin IHC images were analyzed in reference to previous publications(Chang *et al*., 2019; Wu *et al*., 2021a).

#### RCAS virus overexpression

Sprty2 and Wnt3a, overexpression construct, cloning and injection was performed in reference to Wu et al(Wu *et al*., 2018b).

#### Histological preparations, in situ hybridization, RNA scope

The skin explants were fixed in 4% PFA and dehydrated in a graded alcohol series. The tissue was cleared in Xylene and embedded in paraffin wax. 6 μm sections were cut on a microtome. H&E and IHC staining were performed according to accepted protocol. The dilution ratio for section IHC was 1:50. In situ hybridization and RNAscope assays were performed according to Wu et al(Wu *et al*., 2021b).

#### Inhibitor treatment

Blebbistatin (Cayman, MI, USA) and LY2107621 (Cayman, MI, USA) were dissolved in DMSO and added into the medium to reach 10 uM.

#### Phenomenological model

To model the initiation of invagination, a cylindrical coordinate system (*r*, *θ*, *z*) with azimuthal symmetry is applied to the system, with the epidermal layer located at *z* = 0 and the DC located underneath the origin (Fig. 4C, Oblique view). For simplicity, we neglected the proliferation of epidermal cells at this stage, which could potentially lead to epithelial buckling and undulation. Since the model focused on invagination, we also neglected the upward deformation of the epidermal layer caused by the protrusion of the DC. The region with the highest concentration of chemoattractant Tgfβ is represented by a point *Q* located at *z* = − *H*_0_, and the DC is approximated by a cylinder with a radius of *r*_0_. Due to the synergistic effect between MMP and Tgfβ, the spatial profile of MMP activity was assumed to correlate with the profile of Tgfβ. The deformation of epidermal cells at the location *r* is denoted as *u_r_*(*r*) and *u_z_*(*r*) along the radial and the *z* directions, respectively, while *F_c_* represents the chemotactic force acting on the epidermal cells from the dermis (Fig. 4C, Lateral view). The spatial profile of Tgfβ/MMP was approximated by the steady-state solution. The gradient of this solution was used to indicate the force (refer to Equations (S.1-S.3) in the SI). The aspect ratio of the radial length to the azimuthal length of the epidermal cells at the location *r*, denoted as *AS*(*r*), is expressed as follows (refer to Equation (S.4) in the SI),

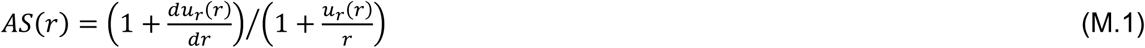

while the changes in cell stiffness were assumed to be linearly proportional to the increase in cell length due to stretching (refer to Equation (S.5) in the SI). The steady-state profiles of *u_r_*(*r*) and *u_z_*(*r*) was numerically estimated using Lamé’s constitutive equation(Sadd, 2014) (refer to Equations (S.9-S.12) in the SI). The results were used to calculate the aspect ratio (Fig. 4E, blue curves) and the changes in cell stiffness (Fig. 4E, green curves). To model the invagination of epidermal cells into the dermis, we considered the combined effect of mechanical resistance from the DC and the mechanical weakening of the basement membrane (BM) by MMP (Fig. 4B). 4B). Due to the diffusion of MMP, epidermal cells on top of the DC encounter direct resistance, whereas epidermal cells at the border of the DC experience a reduction in resistance (Fig. 4B). Only when the force is larger than the resistance, the epidermal cells are allowed to deform in the *z*-direction. Using this approach, we numerically estimated the effective force for the invagination (Fig. 4E, red curve; refer to Equation (S.13) in the SI).

To model the termination of invagination, we tracked the position of leading cells at the forefront of invagination. The position is defined as *r_L_*(*t*) and *z_L_*(*t*) for the radial and *z* coordinates, respectively, while the angular coordinate was ignored due to azimuthal symmetry (Fig. 4D). Since these cells are approaching the region with the highest concentration of Tgfβ, we assumed that the major force acting at these cells is the chemotactic force and the resistance from the dermis (refer to Equations (S.15) and (S.17) in the SI). We also assumed that these cells change stiffness mainly in response to the high concentrations of Tgfβ. For simplicity, we assumed that the increase in cell stiffness of these cells is directly proportional to the concentration of Tgfβ in their immediate vicinity (see Equation (S.2) in the SI). Furthermore, we considered the non-linear effect in the compressive stress-strain relations of the dermis, given that the migratory force from the leading cells is significant. This nonlinear effect indicates an increase in the stiffness of the dermis (refer to Equation (S.16) in the SI), denoted as Δ*λ_D_*:

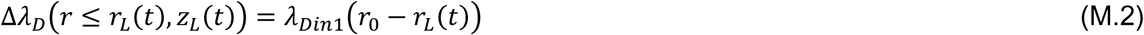

where *λ_Din_*_1_ represents the coefficient of the nonlinear compressive effect of the dermis. Using these approaches, we numerically obtained the path of invagination (Fig. 4F), the changes in the stiffness of the leading cells (Fig. 4G), and the changes in the stiffness of the DC (Fig. 4G) (refer to the SI for more details).

## Supporting information

SI for model detail

SVideo1

SVideo2

SVideo3

SVideo4

SVideo5

## Acknowledgement

This work was supported by National Institutes of Health (NIH) grants R37 AR060306, R35GM153402, RO1 AR047364 and RO1 AR078050 (C.M. Chuong), R01 HL121365 and R35 GM140929 (Y Wang), and the research contract between USC and China Medical University in Taiwan, contract number 005884. WTJ would like to thank services provided by Two-photon imaging facility, China Medical University Hospital, Taiwan. WTJ was supported by the National Science and Technology Council, Taiwan (NSTC 112-2112-M-039-001, NSTC 112-2811-M-039-002), and China Medical University, Taiwan (CMU111-MF-18 and CMU112-MF-01). CLG is funded by NSTC 112-2112-M-001-069, NSTC 112-2314-B-001-001, NSTC 113-2327-B-038-001, AS-GCS-112-M01, AS-GCS-112-M04.

## Author contribution

HIH, TXJ, CHH, JL and PW performed experiments. WTJ, TCC, and WCL performed qMorF analysis. CLG generated the mathematical model. YW provided MMP-FRET materials and analysis guidelines. HIH, PW, WTJ, CJW, CLG and CMC fostered the concepts of the project. HIH, TYL, WTJ, CJW, CLG, CMC contributed to writing the paper.

## SI Fig Legend

**SI Fig 1.**
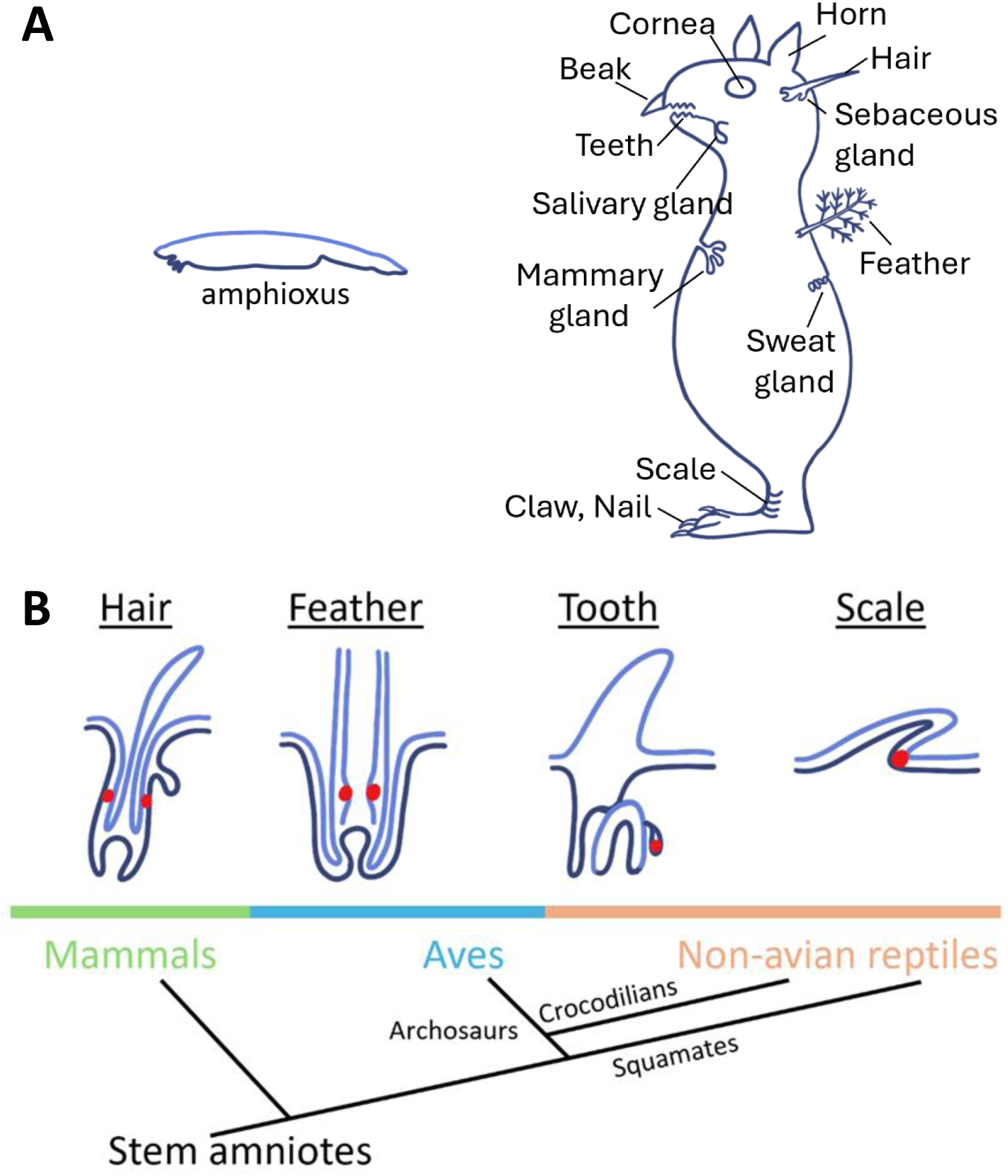
Evo-devo of integumentary organs. A. Ancestral chordates such as amphioxus has smooth integument. As vertebrates evolve, different types of skin appendages emerge to help animals interact with the environment. Shown here is a conceptual animal with different integumentary organs. Modified from Chuong CM edit, 1998. Molecular Basis of Epithelial Appendage Morphogenesis. Landes Biosciences. B. Among these integumentary appendages, the follicle architecture provides epidermal stem cells (red dots) with a dermal niche, facilitating the molting the distal differentiated structures and renewal of new appendages. Follicles in hair, feather and reptile teeth result from convergent evolution. Scales do not have this configuration and keep homeostasis similar to the mechanism used in the epidermis.

**SI Fig 2.**
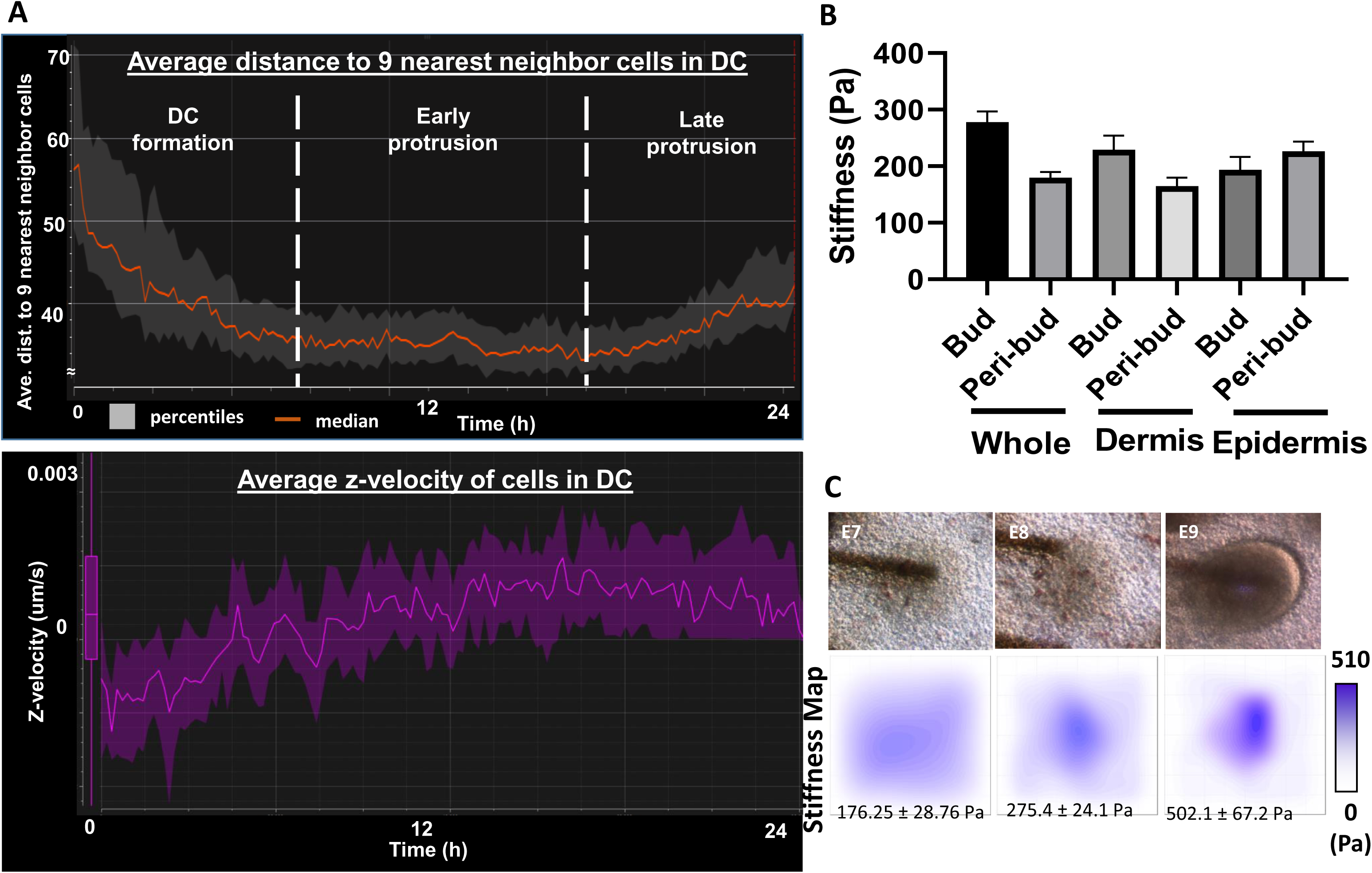
Schematic illustration of feather follicle formation and some biophysical characterization. A. Top panel, Average distance to 9 nearest neighbor cells. Lower panel, average z-velocity in the DC during feather bud protrusion. B. Average tissue stiffness of whole, dermis and epidermis of bud and peri-bud regions of E8 skin. C. Stiffness map of E7, E8 and E9 feather buds showing during dermal condensation and feather bud protrusion, the regional stiffness gradually increases from 176 ± 28 Pa to 275 ± 24 and 502 ± 67 Pa, respectively.

**SI Fig 3.**
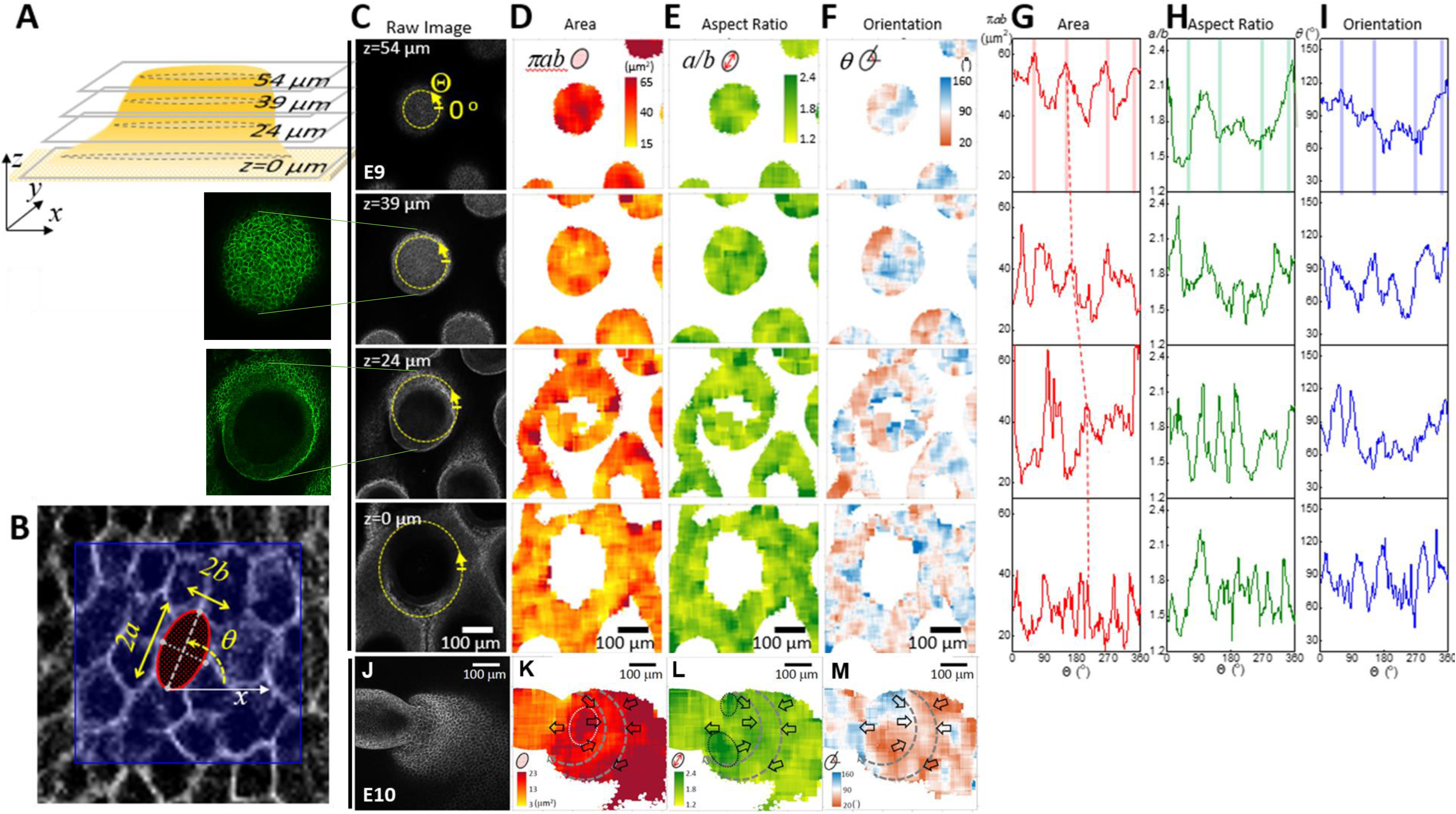
QMorF (Image-cased quantitative morphology field measurement) A. Illustration of different section levels of an E9 feather bud that are used for analyses. B. Representative quantifiable parameters of a cross-sectional cellular morphology in the QMorF analysis. C. Raw image of respective sections. The green lines point to the magnified original E-cadherin staining images of respective feather bud z-sections. D-F. Area, aspect ratio and orientation of respective sections. G-I. Quantified area, aspect ratio and orientation of cell shape over the azimuthal direction (Θ-axis) along the yellow dot circle in panel C of respective sections. Four peaks distributed over the azimuthal direction of the area plot in panel G at the bud tip (z=54 μm) suggest a 4-fold morphological symmetry of cells to accommodate the confided tip geometry. J-M. Whole mount of E10 skin stained with antibodies to L-CAM (E cadherin). Raw image used for quantification of area, aspect ratio and orientation of invaginating E10 feather bud. Please see methods for QMorF analysis.

**SI Fig 4.**
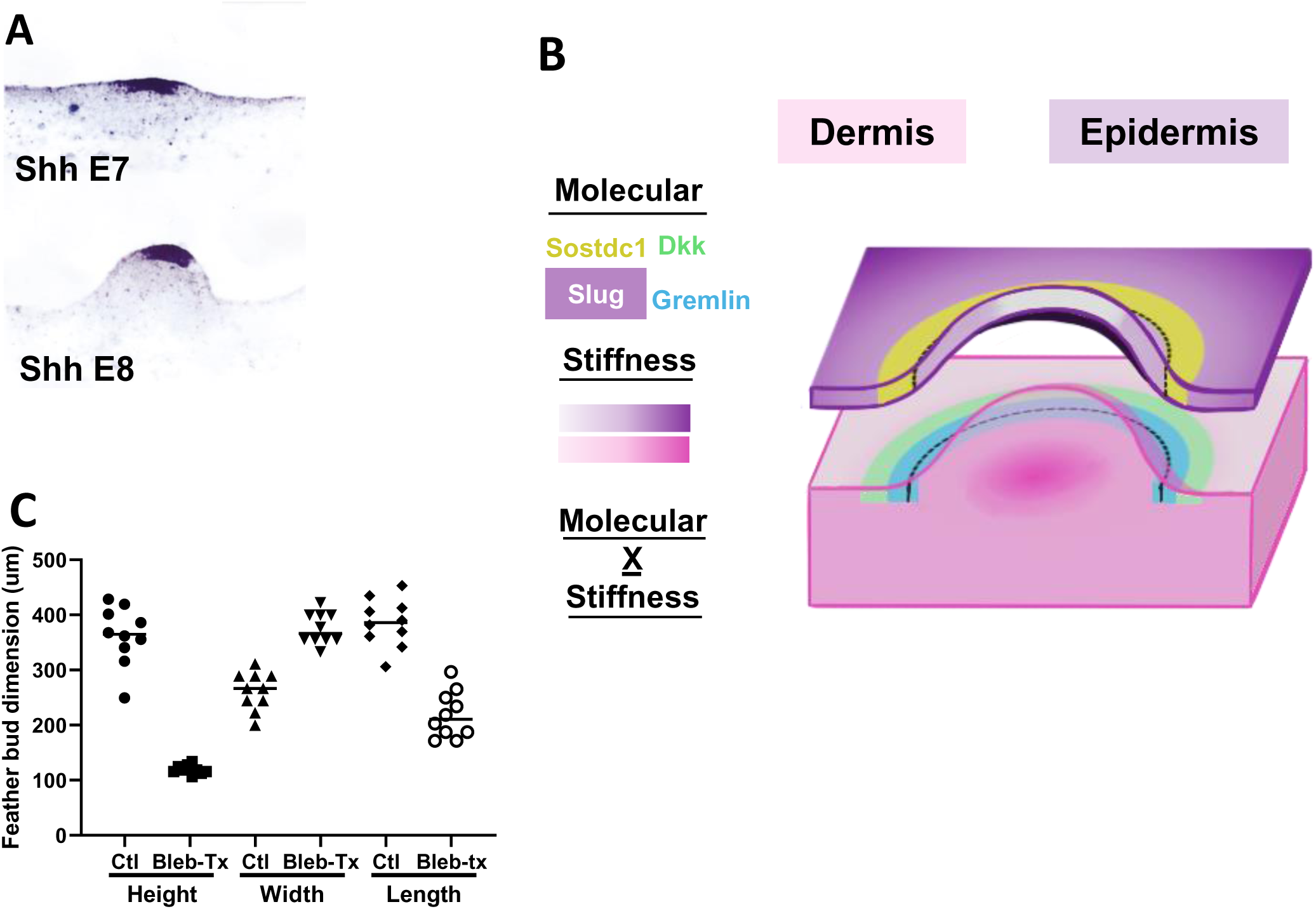
Changes of tissue rigidity and molecular expression during follicle formation. A. In situ hybridization of Shh of E7 and E8 chicken skin B. Spatial expression pattern of Sostdc1, Slug, DKK1 and Gremlin in conjunction with spatial stiffness in the E9 dermis and epidermis. Notice these molecules demarcate bud boundary. C. Quantified dot graph of feather bud dimension changes with and without Bleb-Tx.

**SI Fig 5.**
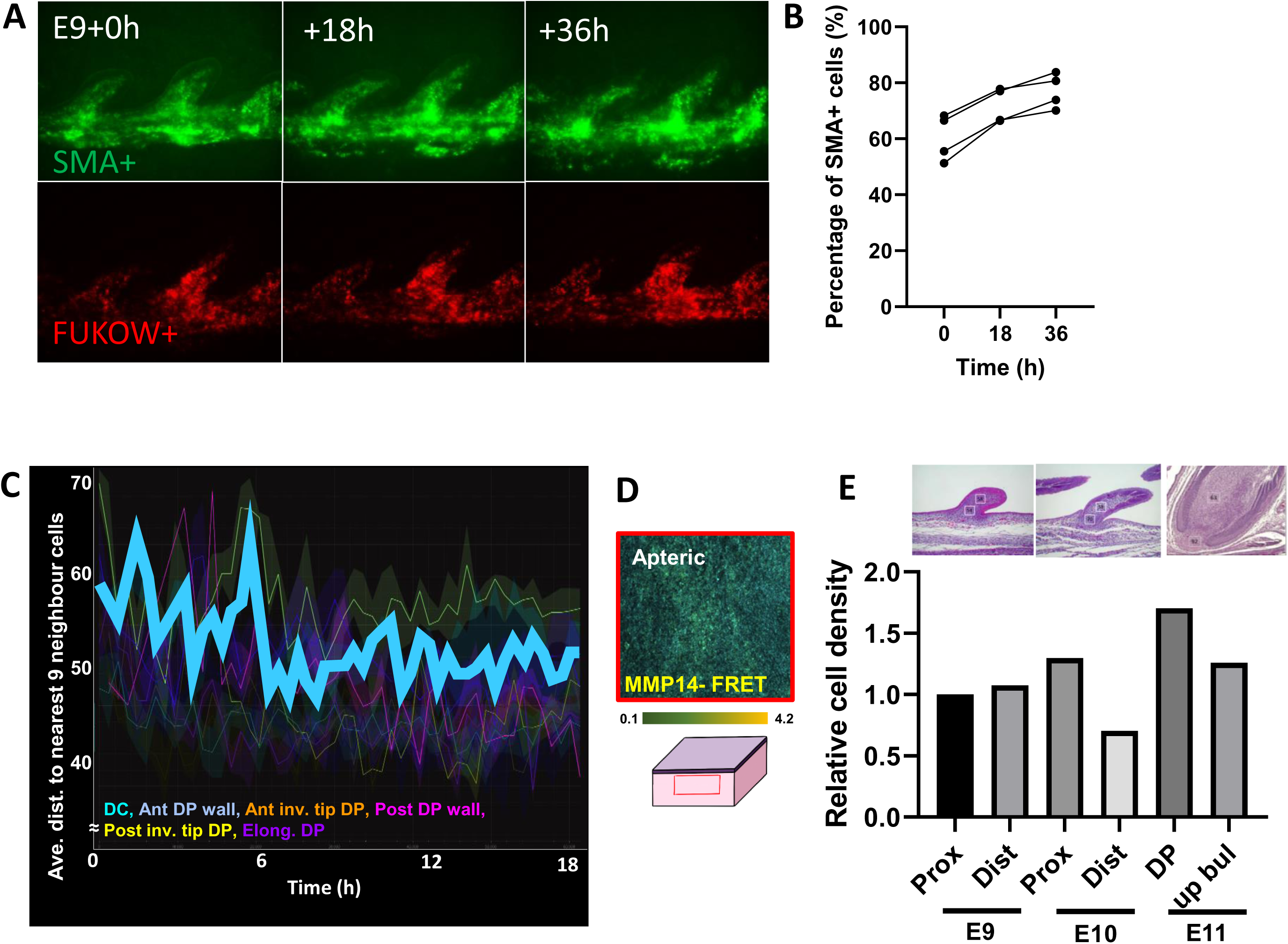
Molecular expression and cellular dynamics of invaginating feather follicle. A. The changes in SMA+ (green) and overall cell (red, FUKOW+) in growing E9 explant for 36h. B. Graph showing quantified percentage of SMA+ over FUKOW+ cells in 4 E9 explant feather follicles. C. Average distance to nearest neighbor cells in the invaginating E11 feather follicle. The bold cyan line indicates the median value of presumptive dermal papilla cells. D. MMP14-FRET activity at the apteric region of E11 chicken skin, as illustrated. E. Relative cell density at proximal, distal, DP or upper bulge regions of the feather bud in E9, E10 and E11 feather bud.

## Reference

1. Bailles A, Collinet C, Philippe JM, Lenne PF, Munro E, Lecuit T (2019) Genetic induction and mechanochemical propagation of a morphogenetic wave. Nature 572: 467–473

2. Bao W, Greenwold MJ, Sawyer RH (2016) Expressed miRNAs target feather related mRNAs involved in cell signaling, cell adhesion and structure during chicken epidermal development. Gene 591: 393–402

3. Campas O, Mahadevan L (2009) Shape and dynamics of tip-growing cells. Curr Biol 19: 2102–2107

4. Chang WL, Wu H, Chiu YK, Wang S, Jiang TX, Luo ZL, Lin YC, Li A, Hsu JT, Huang HL et al (2019) The Making of a Flight Feather: Bio-architectural Principles and Adaptation. Cell 179: 1409–1423 e1417

5. Chen CF, Foley J, Tang PC, Li A, Jiang TX, Wu P, Widelitz RB, Chuong CM (2015) Development, regeneration, and evolution of feathers. Annu Rev Anim Biosci 3: 169–195

6. Chen CK, Chang YM, Jiang TX, Yue Z, Liu TY, Lu J, Yu Z, Lin JJ, Vu TD, Huang TY et al (2024) Conserved regulatory switches for the transition from natal down to juvenile feather in birds. Nat Commun 15: 4174

7. Chuong CM, Chodankar R, Widelitz RB, Jiang TX (2000) Evo-devo of feathers and scales: building complex epithelial appendages. Curr Opin Genet Dev 10: 449–456

8. Chuong CM, Edelman GM (1985) Expression of cell-adhesion molecules in embryonic induction. I. Morphogenesis of nestling feathers. J Cell Biol 101: 1009–1026

9. Collinet C, Lecuit T (2021) Programmed and self-organized flow of information during morphogenesis. Nat Rev Mol Cell Biol 22: 245–265

10. Dhouailly D (2009) A new scenario for the evolutionary origin of hair, feather, and avian scales. J Anat 214: 587–606

11. Dhouailly D (2023) Evo Devo of the Vertebrates Integument. J Dev Biol 11

12. Di-Poi N, Milinkovitch MC (2016) The anatomical placode in reptile scale morphogenesis indicates shared ancestry among skin appendages in amniotes. Sci Adv 2: e1600708

13. Fuchs E (2009) The tortoise and the hair: slow-cycling cells in the stem cell race. Cell 137: 811–819

14. Fuchs E, Merrill BJ, Jamora C, DasGupta R (2001) At the roots of a never-ending cycle. Dev Cell 1: 13–25

15. Gladilin E, Ohse S, Boerries M, Busch H, Xu C, Schneider M, Meister M, Eils R (2019) TGFbeta-induced cytoskeletal remodeling mediates elevation of cell stiffness and invasiveness in NSCLC. Sci Rep 9: 7667

16. Halder G, Dupont S, Piccolo S (2012) Transduction of mechanical and cytoskeletal cues by YAP and TAZ. Nat Rev Mol Cell Biol 13: 591–600

17. Harn HI, Ogawa R, Hsu CK, Hughes MW, Tang MJ, Chuong CM (2019) The tension biology of wound healing. Exp Dermatol 28: 464–471

18. Harn HI, Wang SP, Lai YC, Van Handel B, Liang YC, Tsai S, Schiessl IM, Sarkar A, Xi H, Hughes M et al (2021) Symmetry breaking of tissue mechanics in wound induced hair follicle regeneration of laboratory and spiny mice. Nat Commun 12: 2595

19. Jamora C, DasGupta R, Kocieniewski P, Fuchs E (2003) Links between signal transduction, transcription and adhesion in epithelial bud development. Nature 422: 317–322

20. Jiang T, Secor M, Lansford R, Widelitz RB, Chuong CM (2023) Using Avian Skin Explants to Study Tissue Patterning and Organogenesis. J Vis Exp

21. Jiang TX, Tuan TL, Wu P, Widelitz RB, Chuong CM (2011) From buds to follicles: matrix metalloproteinases in developmental tissue remodeling during feather morphogenesis. Differentiation 81: 307–314

22. Kiat Y, Balaban A, Sapir N, O’Connor JK, Wang M, Xu X (2020) Sequential Molt in a Feathered Dinosaur and Implications for Early Paravian Ecology and Locomotion. Curr Biol 30: 3633–3638 e3632

23. Kim OV, Litvinov RI, Chen J, Chen DZ, Weisel JW, Alber MS (2017) Compression-induced structural and mechanical changes of fibrin-collagen composites. Matrix Biol 60-61: 141–156

24. Kobayashi T, Kim H, Liu X, Sugiura H, Kohyama T, Fang Q, Wen FQ, Abe S, Wang X, Atkinson JJ et al (2014) Matrix metalloproteinase-9 activates TGF-beta and stimulates fibroblast contraction of collagen gels. Am J Physiol Lung Cell Mol Physiol 306: L1006–1015

25. Lai YC, Liang YC, Jiang TX, Widelitz RB, Wu P, Chuong CM (2018) Transcriptome analyses of reprogrammed feather / scale chimeric explants revealed co-expressed epithelial gene networks during organ specification. BMC Genomics 19: 780

26. Leggett SE, Hruska AM, Guo M, Wong IY (2021) The epithelial-mesenchymal transition and the cytoskeleton in bioengineered systems. Cell Commun Signal 19: 32

27. Lei M, Harn HI, Li Q, Jiang J, Wu W, Zhou W, Jiang TX, Wang M, Zhang J, Lai YC et al (2023) The mechano-chemical circuit drives skin organoid self-organization. Proc Natl Acad Sci U S A 120: e2221982120

28. Lenne PF, Trivedi V (2022) Sculpting tissues by phase transitions. Nat Commun 13: 664

29. Li A, Chen M, Jiang TX, Wu P, Nie Q, Widelitz R, Chuong CM (2013) Shaping organs by a wingless-int/Notch/nonmuscle myosin module which orients feather bud elongation. Proc Natl Acad Sci U S A 110: E1452–1461

30. Li A, Cho JH, Reid B, Tseng CC, He L, Tan P, Yeh CY, Wu P, Li Y, Widelitz RB et al (2018) Calcium oscillations coordinate feather mesenchymal cell movement by SHH dependent modulation of gap junction networks. Nat Commun 9: 5377

31. Li A, Figueroa S, Jiang TX, Wu P, Widelitz R, Nie Q, Chuong CM (2017) Diverse feather shape evolution enabled by coupling anisotropic signalling modules with self-organizing branching programme. Nat Commun 8: ncomms14139

32. Li KN, Tumbar T (2021) Hair follicle stem cells as a skin-organizing signaling center during adult homeostasis. EMBO J 40: e107135

33. Martino P, Sunkara R, Heitman N, Rangl M, Brown A, Saxena N, Grisanti L, Kohan D, Yanagisawa M, Rendl M (2023) Progenitor-derived endothelin controls dermal sheath contraction for hair follicle regression. Nat Cell Biol 25: 222–234

34. Massague J, Sheppard D (2023) TGF-beta signaling in health and disease. Cell 186: 4007–4037

35. Melisi D, Ishiyama S, Sclabas GM, Fleming JB, Xia Q, Tortora G, Abbruzzese JL, Chiao PJ (2008) LY2109761, a novel transforming growth factor beta receptor type I and type II dual inhibitor, as a therapeutic approach to suppressing pancreatic cancer metastasis. Mol Cancer Ther 7: 829–840

36. Mongera A, Rowghanian P, Gustafson HJ, Shelton E, Kealhofer DA, Carn EK, Serwane F, Lucio AA, Giammona J, Campas O (2018) A fluid-to-solid jamming transition underlies vertebrate body axis elongation. Nature 561: 401–405

37. Moore-Smith LD, Isayeva T, Lee JH, Frost A, Ponnazhagan S (2017) Silencing of TGF-beta1 in tumor cells impacts MMP-9 in tumor microenvironment. Sci Rep 7: 8678

38. Morita R, Fujiwara H (2022) Tracing the developmental origin of tissue stem cells. Dev Growth Differ 64: 566–576

39. Morita R, Sanzen N, Sasaki H, Hayashi T, Umeda M, Yoshimura M, Yamamoto T, Shibata T, Abe T, Kiyonari H et al (2021) Tracing the origin of hair follicle stem cells. Nature 594: 547–552

40. Morrison SJ, Spradling AC (2008) Stem cells and niches: mechanisms that promote stem cell maintenance throughout life. Cell 132: 598–611

41. Panciera T, Azzolin L, Cordenonsi M, Piccolo S (2017) Mechanobiology of YAP and TAZ in physiology and disease. Nat Rev Mol Cell Biol 18: 758–770

42. Prum RO (2005) Evolution of the morphological innovations of feathers. J Exp Zool B Mol Dev Evol 304: 570–579

43. Prum RO (2010) Moulting tail feathers in a juvenile oviraptorisaur. Nature 468: E1; discussion E2

44. Sadd MH (2014) Elasticity. In: Elasticity, p. 103. Academic Press: Oxford

45. Schneider AK, Cama G, Ghuman M, Hughes FJ, Gharibi B (2017) Sprouty 2, an Early Response Gene Regulator of FosB and Mesenchymal Stem Cell Proliferation During Mechanical Loading and Osteogenic Differentiation. J Cell Biochem 118: 2606–2614

46. Shintani Y, Fukumoto Y, Chaika N, Grandgenett PM, Hollingsworth MA, Wheelock MJ, Johnson KR (2008) ADH-1 suppresses N-cadherin-dependent pancreatic cancer progression. Int J Cancer 122: 71–77

47. Song H, Wang Y, Goetinck PF (1996) Fibroblast growth factor 2 can replace ectodermal signaling for feather development. Proc Natl Acad Sci U S A 93: 10246–10249

48. Spanjer AI, Baarsma HA, Oostenbrink LM, Jansen SR, Kuipers CC, Lindner M, Postma DS, Meurs H, Heijink IH, Gosens R et al (2016) TGF-beta-induced profibrotic signaling is regulated in part by the WNT receptor Frizzled-8. FASEB J 30: 1823–1835

49. Stenn KS, Paus R (2001) Controls of hair follicle cycling. Physiol Rev 81: 449–494

50. Sternlicht MD, Kouros-Mehr H, Lu P, Werb Z (2006) Hormonal and local control of mammary branching morphogenesis. Differentiation 74: 365–381

51. Strzyz P (2019) Forcing through barriers. Nat Rev Mol Cell Biol 20: 136

52. Ting-Berreth SA, Chuong CM (1996) Local delivery of TGF beta2 can substitute for placode epithelium to induce mesenchymal condensation during skin appendage morphogenesis. Dev Biol 179: 347–359

53. Tozluoglu M, Duda M, Kirkland NJ, Barrientos R, Burden JJ, Munoz JJ, Mao Y (2019) Planar Differential Growth Rates Initiate Precise Fold Positions in Complex Epithelia. Dev Cell 51: 299–312 e294

54. Tronci G, Doyle A, Russell SJ, Wood DJ (2013) Triple-helical collagen hydrogels via covalent aromatic functionalization with 1,3-Phenylenediacetic acid. J Mater Chem B 1: 5478–5488

55. Vinciguerra P, Mencucci R, Romano V, Spoerl E, Camesasca FI, Favuzza E, Azzolini C, Mastropasqua R, Vinciguerra R (2014) Imaging mass spectrometry by matrix-assisted laser desorption/ionization and stress-strain measurements in iontophoresis transepithelial corneal collagen cross-linking. Biomed Res Int 2014: 404587

56. Wang X, O’Connor J, Zheng X, Wang Y, Kiat Y (2024) Earliest evidence of avian primary feather moult. Biol Lett 20: 20240106

57. Widelitz RB, Jiang TX, Lu J, Chuong CM (2000) beta-catenin in epithelial morphogenesis: conversion of part of avian foot scales into feather buds with a mutated beta-catenin. Dev Biol 219: 98–114

58. Widelitz RB, Jiang TX, Noveen A, Chen CW, Chuong CM (1996) FGF induces new feather buds from developing avian skin. J Invest Dermatol 107: 797–803

59. Wollensak G, Spoerl E, Seiler T (2003) Stress-strain measurements of human and porcine corneas after riboflavin-ultraviolet-A-induced cross-linking. J Cataract Refract Surg 29: 1780–1785

60. Wu H, Chiu YK, Tsai JC, Chuong CM, Juan WT (2021a) A quantitative image-based protocol for morphological characterization of cellular solids in feather shafts. STAR Protoc 2: 100661

61. Wu P, Hou L, Plikus M, Hughes M, Scehnet J, Suksaweang S, Widelitz R, Jiang TX, Chuong CM (2004) Evo-Devo of amniote integuments and appendages. Int J Dev Biol 48: 249–270

62. Wu P, Jiang TX, Lei M, Chen CK, Hsieh Li SM, Widelitz RB, Chuong CM (2021b) Cyclic growth of dermal papilla and regeneration of follicular mesenchymal components during feather cycling. Development 148

63. Wu P, Lai YC, Widelitz R, Chuong CM (2018a) Comprehensive molecular and cellular studies suggest avian scutate scales are secondarily derived from feathers, and more distant from reptilian scales. Sci Rep 8: 16766

64. Wu P, Wu X, Jiang TX, Elsey RM, Temple BL, Divers SJ, Glenn TC, Yuan K, Chen MH, Widelitz RB et al (2013) Specialized stem cell niche enables repetitive renewal of alligator teeth. Proc Natl Acad Sci U S A 110: E2009–2018

65. Wu P, Yan J, Lai YC, Ng CS, Li A, Jiang X, Elsey RM, Widelitz R, Bajpai R, Li WH et al (2018b) Multiple Regulatory Modules Are Required for Scale-to-Feather Conversion. Mol Biol Evol 35: 417–430

66. Wu XS, Yeh CY, Harn HI, Jiang TX, Wu P, Widelitz RB, Baker RE, Chuong CM (2019) Self-assembly of biological networks via adaptive patterning revealed by avian intradermal muscle network formation. Proc Natl Acad Sci U S A 116: 10858–10867

67. Xu X, Zheng X, You H (2010) Exceptional dinosaur fossils show ontogenetic development of early feathers. Nature 464: 1338–1341

68. Xu X, Zhou Z, Dudley R, Mackem S, Chuong CM, Erickson GM, Varricchio DJ (2014) An integrative approach to understanding bird origins. Science 346: 1253293

69. Yang S, Palmquist KH, Nathan L, Pfeifer CR, Schultheiss PJ, Sharma A, Kam LC, Miller PW, Shyer AE, Rodrigues AR (2023) Morphogens enable interacting supracellular phases that generate organ architecture. Science 382: eadg5579

70. Yu M, Wu P, Widelitz RB, Chuong CM (2002) The morphogenesis of feathers. Nature 420: 308–312

71. Yue Z, Jiang TX, Widelitz RB, Chuong CM (2005) Mapping stem cell activities in the feather follicle. Nature 438: 1026–1029

72. Yue Z, Jiang TX, Wu P, Widelitz RB, Chuong CM (2012) Sprouty/FGF signaling regulates the proximal-distal feather morphology and the size of dermal papillae. Dev Biol 372: 45–54

