## Supplementary material for "Novel Tissue Mechanics-Guided Cellular Flows Enable the Evolution of Feather Follicles": SI for model detail

### Long Version for Supplemental Information

#### The Phenomenological Chemotactic Model for Epidermal Invagination

#### 1. The Assumptions and Aims of the Model

Upon the condensation of dermal cells, a switch in dermal cell movement was observed, correlating with alterations in the shape and rigidity of epidermal cells. Subsequently, epidermal invagination occurred around the dermal condensate (DC), halting once the invagination reached the base of the DC. Conceptualizing the FGF-stimulated DC cells as confined hot gas molecules by the epidermis, a probable mechanism for initiating their vertical upward motion is to establish a mechanically weak point in the epidermis above the DC. How could the weakening of epidermis be initiated and confined at the region above the DC? Mechanically, condensed dermal cells at the DC apply traction forces on the neighboring tissue. While horizontally or vertically downward forces can be countered by traction forces from surrounding DCs or by dermal resistance, vertically upward forces eventually reach the epidermal layer, exerting downward pulling forces on epidermal cells. Additionally, dermal cell-secreted chemical factors like Tgfβ can stimulate epidermal cells, prompting morphogenetic movements. Tgfβ can also signal surrounding dermal cells to increase MMP secretion[1], further activating Tgfβ[2]. As MMP can degrade the basement membrane (BM)[3], the combined effect of Tgfβ and MMP promotes the formation of weakened regions in the BM near the DC. The first aim of our modeling is to explore whether the mechano-chemical activities of dermal cells can explain the observed changes in epidermal cell shape and stiffness during early bud protrusion, particularly the weak point formation. Conversely, if Tgfβ-MMP signaling triggers epidermal cell migration, it is probable that migrating epidermal cells with higher stiffness tend to invade softer regions rather than stiffer ones. The diffusivity of Tgfβ and MMP extends the weakening effect around the DC periphery, where migratory epidermal cells encounter less mechanical resistance. The second aim of our modeling is to assess whether the combination of DC stiffness and Tgfβ-MMP signaling aids in specifying the location of epidermal invagination. The area with the highest Tgfβ-MMP signaling activity likely resides near the base of the DC, prompting epidermal cells to move towards this target through invagination. Throughout invagination, both epidermal and dermal cells alter their stiffness due to their mechano-chemical interactions, consequently modifying the invagination pattern of migratory epidermal cells. The third aim of our modeling is to explore whether this interplay can determine the termination of invagination.

#### 2. The Initiation of Invagination

In this section, we focused on the initiation stage right before the onset of invagination. A coarse-grained phenomenological model was developed to investigate the force distribution and deformation of epidermal cells. In this stage, the cross-sectional view of the follicle revealed a deformation of the epidermis directly above the DC, corresponding to the placode (Fig. 1A, E7.5), which eventually develops into the distal bud end. The geometric influence of this region complicates the analysis. Moreover, we considered that the changes in cell area and aspect ratio at the distal bud end are primarily due to the physical protrusion from the DC/CP (Fig. 1F). Since the current model focused on the Tgfβ-MMP effect, we disregarded the distal bud end region and treated the entire epidermal layer as a flat, planar sheet. The traction force from the DC can be highly dynamic and heterogeneous. For simplicity, we only considered the impact of Tgfβ-MMP signaling on the epidermal cells. We further approximated the DC as a cylindrical structure with a radius of $r_0$ (Fig. 4D). Due to the specificity of cell-cell adhesions, the mixing of epidermal cells with dermal cells is prohibited. In this context, we treated the epidermal layer and the DC as two separate entities with continuous borders of their own. The invagination of the epidermal layer around the DC is considered a spatial competition process. Due to the dense packing of dermal cells in the condensate, the epidermal cells can push the DC inward only if their migratory force is sufficiently large. A cylindrical coordinate system ($r$, $\theta$, $z$) is used to describe the dynamics of the epidermal layer. The epidermal layer is positioned at the plane of $z$ equal to zero, and the projected point of the center of the DC cylinder onto the epidermal layer serves as the origin. Azimuthal undulation was also disregarded in the current model. This simplification allows us to impose azimuthal symmetry on the epidermal layer with respect to the center and ignore the variations along the $\theta$ direction.

To model epidermal invagination, we assumed that the movement of epidermal cells is a chemotaxis-like response to diffusive Tgfβ-MMP signaling produced by the dermal cells. Such a response often enhances the polymerization and reorganization of the cytoskeleton, which, in turn, can increase cell stiffness. For simplicity, we assumed that the magnitude of migratory force and the increment of cell stiffness of epidermal cells are proportional to the Tgfβ-MMP signaling. Since cells often adapt to sustained activation, we approximated the effective Tgfβ-MMP signaling as the gradient of a diffusive activity. From the experimental data, it is evident that the movement and proliferation of dermal cells are highly dynamic and heterogeneous during follicle formation. The activity of these cells in secreting and activating Tgfβ and MMP can vary significantly. To simplify, we approximated the Tgfβ-MMP signaling as being generated from a point source, $Q$, in the central area of the condensate cylinder. The distance from $Q$ to the epidermal layer is denoted as "$H_0$." The strength of the Tgfβ-MMP signaling is denoted as "$C$." The spatial profile of $C$ follows:

$$\frac{dC}{dt} = -k_C C + D_C \nabla^2 C \tag{S.1}$$

In Equation (S.1), $k_C$ and $D_C$ are the adaptation rate and diffusion constant of the Tgfβ-MMP signaling, respectively. Since biomolecules typically diffuse faster than cell migration, we adopted a simple approach by assuming that $C$ always reaches its steady state during follicle formation. In 3D space, the solution to equation (1) is as follows:

$$C(R) = C_0\, e^{-R/\sqrt{D_C/k_C}}/R \tag{S.2}$$

where $C_0$ is a constant and $R$ represents the distance from the epidermal cells of interest to the point $Q$. For convenience, we set the diffusion length as $l = (D_C/k_C)^{1/2}$. The chemotactic force acting on the epidermal cells was approximated as:

$$\boldsymbol{F}_C(R) = \nabla C(R) = -C(R)(l^{-1} + R^{-1})\,\boldsymbol{R}/R \tag{S.3}$$

where we used bold text to indicate the vector form of a symbol, e.g., $\boldsymbol{R} \stackrel{\text{def}}{=} \vec{R}$. Conversely, a symbol not in bold text represents the scalar form of the vector or tensor, for example, $R = |\boldsymbol{R}|$. The Tgfβ-MMP signaling guides the epidermal cells to invaginate and approach the point source $Q$. From

Equation (S.3), we noted that the force increases as the distance $R$ decreases. Consequently, when the invagination of epidermal cells is deep enough to reach the point source $Q$, the magnitude of migratory force can be sufficiently large to overcome the resistance from the DC and initiate centripetal movement of epidermal cells. This movement, in turn, pushes the DC inward and facilitates the closure of the follicle base.

In addition to the Tgfβ-MMP signaling, another factor that influences the deformation and displacement of the epidermal layer is the epidermal elastic energy. For simplicity, we used Lame's constitutive equation[4] to approximate the elastic responses of the epidermal layer. The displacement of the epidermal layer at the position $(r, \theta, 0)$ is denoted as $\boldsymbol{u}(r, \theta, 0)$. With the azimuthal symmetry of the epidermal layer and the fact that the epidermal layer is positioned at a constant plane of $z$ equal to zero, we defined $\boldsymbol{u}(r, \theta, 0)$ as $\boldsymbol{u}(r)$. Moreover, we separated $\boldsymbol{u}(r)$ into the in-plane (tangential to the epidermal layer) component, $\boldsymbol{u_t}(r) = \boldsymbol{u_r}(r)$, and the out-of-plane (normal to the epidermal layer) component, $\boldsymbol{u_n}(r) = \boldsymbol{u_z}(r)$. Here, "$\boldsymbol{u_z}$" represents the chemotaxis-induced morphological changes or invasion of epidermal cells. This component is characterized by the extension of the cell body or the formation of cell protrusions into the dermis. In comparison, "$\boldsymbol{u_r}$" represents the shift of the cell boundary in the radial direction. According to the strain-displacement relations[5], the change in cell length along the radial direction is indicated by the radial strain $\varepsilon_r = du_r/dr$, whereas the change in cell length along the azimuthal direction is represented by the circumferential strain $\varepsilon_\theta = u_r/r$. Note that the azimuthal components in □□ were neglected due to the azimuthal symmetry in our model. Consequently, the aspect ratio of the radial length to the azimuthal length of the epidermal cells, denoted as $AS(r)$, is expressed as follows:

$$AS(r) = \left(1 + \frac{du_r(r)}{dr}\right) / \left(1 + \frac{u_r(r)}{r}\right) \tag{S.4}$$

In vivo, epithelial cells form a continuous cell sheet that serves as a protective barrier for the organs and the organisms against injuries. This sheet ensures mechanical continuity by organizing actin belts at the cell-cell junctions of each cell, creating a tension network. Stretching epidermal cells within the sheet is likely to increase tension, thereby enhancing cell stiffness to maintain the sheet's integrity. Conversely, compressing cells is unlikely to influence the sheet's integrity and might have minimal effects on cell stiffness. Indeed, it has been shown that stretching epidermal layers can increase cell stiffness, promoting mechanical responses, such as cell proliferation and YAP nuclear translocation[6,7]. Various studies on cell mechanics have also shown that elongating a cell can significantly affect cytoskeletal organization, leading to increased cell stiffness[8-11]. By contrast, the effects of reducing cell length or compressing the cell body on cell stiffness remain unclear. Nevertheless, it is important to note that mechanisms beyond mechanical stretching, such as chemical stimulation, can also enhance cell stiffness. For example, exposing cells to cytokines, such as Tgfβ, can induce cytoskeletal reorganization and increase cell stiffness[12]. This effect can occur even without chemoattractant gradients and often involves YAP signaling[13]. In addition, we should point out that the responses to mechanical compression in the dermis might differ significantly from those in the epidermal layers. Unlike the epidermal layers, which are primarily composed of cells, the dermis contains a substantial amount of extracellular matrix (ECM) molecules, particularly type I collagen fibers, which significantly influence dermal mechanical properties. Studies have shown that compression leads to a nonlinear increase in the stress-strain relations of collagen-based hydrogels[14-17], suggesting that compression can enhance the stiffness of collagen-based matrices such as the dermis.

During the intermediate stage, epidermal cells are positioned at a distance from the source of Tgfβ-MMP signaling. We thus assumed that the observed changes in cell stiffness in these epidermal cells are primarily due to cell elongation or the development of cell protrusions, which subsequently induce cytoskeletal reorganization resulting in increased cell stiffness. To simplify the analysis, we further hypothesized that the rise in cell stiffness is directly proportional to changes in cell length and can be estimated by the cumulative impact of cell stretching in all orientations. A mathematical expression, labeled as $\Delta\lambda(r)$, is introduced to depict the elevation in cell stiffness within the epidermal layers:

$$\Delta\lambda(r) = \Delta\lambda_r \Phi\left(\frac{du_r(r)}{dr}\right) + \Delta\lambda_\theta \Phi\left(\frac{u_r(r)}{r}\right) + \Delta\lambda_z \Phi\big(u_z(r)\big) \quad \text{(S.5)}$$

where $\Phi(x)$ represents a hemi-side function:

$$\Phi(x) = \begin{cases} 0, & x < 0 \\ x, & x \geq 0 \end{cases} \quad \text{(S.6)}$$

In Equation (S.5), "$\lambda$" represents the elastic constant of epidermal cells, also known as Lame's constant. $\lambda$ is often used as a measurement of cell stiffness. In this regard, the coefficients $\Delta\lambda_r$, $\Delta\lambda_\theta$, and $\Delta\lambda_z$ in Equation (S.5) indicate the contribution to the increase in cell stiffness due to stretching in the radial, azimuthal, and vertical directions, respectively.

The experimental findings showed changes in epidermal aspect ratio and cell stiffness as follicle formation progressed. To obtain compatible profiles in the model, a steady-state approach was employed. Cell proliferation within the epidermal layers was not considered for analytical convenience. Furthermore, we assumed that the increase in cell stiffness due to epidermal cell elongation was negligible compared to the zero-order elastic modulus of the cells. This strategy enables the use of a linear approximation to derive the steady-state solution for the equation. Similar to the separation of $\boldsymbol{u}$, we divided the chemotactic force into two components, $\boldsymbol{F_r}$ and $\boldsymbol{F_z}$, as follows:

$$\boldsymbol{F_r}(r) = F_C(R)\,\boldsymbol{r}/R \quad \text{(S.7)}$$

$$\boldsymbol{F_z}(r) = F_C(R) H_0\, \hat{\boldsymbol{z}}/R \quad \text{(S.8)}$$

Next, we sought the equilibrium profiles of displacement $\boldsymbol{u}$. For $\boldsymbol{u_r}$, we assumed that its equation of motion follows Lame's constitutive equation[4]:

$$\xi \frac{d\boldsymbol{u_r}(r)}{dt} = \mu\nabla^2\boldsymbol{u_r}(r) + (\lambda+\mu)\nabla\big(\nabla\cdot\boldsymbol{u_r}(r)\big) + \boldsymbol{F_r}(r) + \nabla\cdot\boldsymbol{T_r} + O(\boldsymbol{u}^2) \quad \text{(S.9)}$$

where $\xi$ represents the viscosity of the epidermal layer, $\mu$ represents the zero-order shear modulus or modulus of rigidity of the epidermal layer, $\lambda$ is the zero-order elastic constant of the epidermal layers, and $\boldsymbol{T_r}$ represents the boundary force tensor for proper boundary conditions at the inter-bud region. In Equation (S.9), the term $O(\boldsymbol{u}^2)$ represents the high-order nonlinear effect that we ignored when solving for the steady-state solution. Setting $r = r_b$ as the boundary at the inter-bud region and $u_r(0) = 0$ at the origin, we have the steady-state solution of $u_r$ as follows:

$$u_r(r) = -r^{\frac{\lambda+\mu}{\lambda+2\mu}} \int_0^r \frac{dx}{x^{\frac{2\lambda+3\mu}{\lambda+2\mu}}} \int_0^x \frac{y dy}{\lambda+2\mu} F_r(y) - \frac{T_r}{\mu} r \tag{S.10}$$

where $T_r$ is the scalar form that can be determined using the boundary condition. Setting $u_r(r_b) = 0$ at the inter-bud region, we have $T_r$ as follows:

$$T_r = -\mu r_b^{-\frac{\mu}{\lambda+2\mu}} \int_0^{r_b} \frac{dx}{x^{\frac{2\lambda+3\mu}{\lambda+2\mu}}} \int_0^x \frac{y dy}{\lambda+2\mu} F_r(y) \tag{S.11}$$

For $\boldsymbol{u}_z$, on the other hand, we assumed that its equation of motion follows a simple elastic response of the dermis before the actual invagination occurs. In other words, we neglected the nonlinear effect in the compressive stress-strain relations of the dermis in the initial phase of invagination, as the displacement is small enough to disregard the nonlinear effect. The equation reads as follows:

$$\xi \frac{d\boldsymbol{u}_z(r)}{dt} = -\lambda_D(r)\boldsymbol{u}_z(r) - \hat{\boldsymbol{z}}\Phi\left(-\left(F_z(r) + B(r)\right)\right) + O(\boldsymbol{u}^2) \tag{S.12}$$

Similar to the concept described in Equation (S.9), the term $O(\boldsymbol{u}^2)$ in Equation (S.12) signifies the high-order nonlinear impact that was disregarded when solving the steady-state solution. Here, $\lambda_D(r)$ denotes the elastic constant of the dermis, which varies with position, while $B(r)$ indicates a position-dependent barrier function that hinders epidermal cell invasion into the dermis. When $r \leq r_0$, the epidermal cells are positioned on top of the DC and encounter greater resistance from the dermis during invagination. When $r > r_0$, on the other hand, the epidermal cells face less resistance from the dermis. To simplify the analysis, we set $B = B_{in}$ and $\lambda_D = \lambda_{Din0}$ for $r \leq r_0$, and $B = B_{out}$ and $\lambda_D = \lambda_{Dout}$ for $r > r_0$. The function $\Phi(x)$ in Equation (15) corresponds to the hemi-side function introduced in Equation (6). This function characterizes the effective force in the $z$ direction. The presence of the barrier modifies the effective force for the epidermal cells to invaginate, $\boldsymbol{F}_{z,eff}(r)$, as:

$$\boldsymbol{F}_{z,eff}(r) = -\hat{\boldsymbol{z}}\Phi\left(-\left(F_z(r) + B(r)\right)\right) \tag{S.13}$$

At the steady state, the displacement in the vertical direction is solved as:

$$u_z(r) = F_{z,eff}(r)/\lambda_D(r) \tag{S.14}$$

Figure 4 B-C and SI Figure 4 A-B show the numerical results. In the simulation, we used $l = 3$, $r_0 = 1$, $\lambda = 0.25$, $\mu = 0.25$, $\lambda_D = \lambda_{Din0} = 100$ for $r \leq r_0$ and $\lambda_D = \lambda_{Dout} = 0.1$ for $r > r_0$, $B = B_{in} = 11$ for $r \leq r_0$ and $B = B_{out} = 0.65$ for $r > r_0$, $C_0 = 20$, and $H_0 = 4$. For the effective force, we considered the dermal resistance and used Equation (S.13) to compute the force.

### 3. The Termination of Invagination

To model the closure of epidermal invagination, we took into account the non-linear effect in the compressive stress-strain relations of the dermis, as evidenced by the deformation of the DC in experimental observations. At this stage, the proliferation of epidermal cells becomes significant and can notably affect the elastic responses of the epidermal layers. To simplify the analysis, our focus was directed towards the movement of leading cells located at the forefront of the invagination. We assumed that a cluster of leading cells positioned at the foremost edge, referred to as the "tongue," initiates the invagination process. The moving trajectories of these cells, denoted as $(r_L(t), \theta, z_L(t))$, with $\theta$ ranging from 0 to $2\pi$ to account for azimuthal symmetry of the invagination, were used to depict the progression of follicle wall development and the closure of the invagination at the base of the DC (Fig. 4D). Given that the invagination brings the leading cells in close proximity to the sources of Tgfβ-MMP signaling, the impact of Tgfβ-MMP signaling is deemed more significant than the elastic responses of the epidermal cells at these leading cells. Consequently, the contribution of the elastic responses of the epidermal layers could be disregarded when analyzing the movement of the leading cells. Likewise, the chemical influence of Tgfβ-MMP signaling on the increase in cell stiffness at the leading cells is considered more substantial compared to the effect of cell stretching. It is thus reasonable to postulate that the chemical effect plays a predominant role in enhancing cell stiffness at the leading cells. For the sake of simplicity, we assumed that the rise in cell stiffness of the leading cells is directly proportional to the activity of Tgfβ-MMP signaling in their immediate vicinity (Equation (S.2)). The initial positions of the leading cells were set at $r_L(0) = r_0$ and $z_L(0) = 0$. Under the assumption that the directional movement of the leading cells adheres to azimuthal symmetry, their radial motion is described by the following equation:

$$\xi \frac{dr_L}{dt} = [\lambda_{Din0} + \lambda_{Din1}(r_0 - r_L)](r_0 - r_L) - \Phi\big(-(F_r(r_L, z_L) + B_{in} - \sigma/r_L)\big) \tag{S.15}$$

In Equation (S.15), the parameter $\lambda_{Din0}$ denotes the zero-order elastic constant of the dermis, while $\lambda_{Din1}$ signifies the coefficient pertaining to the nonlinear compressive effect of the dermis. The hemi-side function $\Phi(\ldots)$ denotes the effective force exerted in the radial direction. This force contains three components: the force $F_r(r_L, z_L)$ derived from the gradients of Tgfβ-MMP signaling, the dermal resistance "$B$", and the tension "$\sigma$"□arising from the actin cable network within the epidermal layer. The "−" sign preceding $\sigma$ indicates that the tension acts inwardly to compress the DC, while the factor "$1/r_L$\" represents the curvature of the epidermal layer encircling the follicle wall. Given that the invagination consistently occurs along the sidewall of the DC, it follows that $r_L \leq r_0$. As such, we used $\lambda_{Din0}$ and $B_{in}$ to represent the zero-order elastic constant and the resistance of the dermis in the radial direction, respectively. In terms of the nonlinear compressive effect, the enhancement in dermal stiffness is expressed as $\lambda_{Din1}(r_0 - r_L)$. To reduce the complexity, we have neglected the contribution from the elastic responses of the epidermal layers in Equation (S.15). Throughout the process of invagination, the DC situated within a radius of $r \leq r_L(t)$ and at the coordinate $z = z_L(t)$ undergoes an increment in its stiffness, denoted as $\Delta\lambda_D$:

$$\Delta\lambda_D\big(r \leq r_L(t), z_L(t)\big) = \lambda_{Din1}\big(r_0 - r_L(t)\big) \tag{S.16}$$

To address the movement of leading cells in the vertical ($z$) direction, $z_L(t)$, we modified Equation (S.12) accordingly. Since the invagination primarily takes place along the sidewall of the DC, the resistance faced by the leading cells in the $z$ direction was approximated using $B_{out}$. Moreover, we assumed that the leading cells actively remodel the dermis to facilitate the invagination process. Consequently, the elastic response component in Equation (S.12) was disregarded when characterizing the movement of leading cells in the $z$ direction. The resulting equation of motion for $z_L$ can be expressed as follows:

$$\xi \frac{dz_L}{dt} = -\Phi\left(-\left(F_z(r_L, z_L) + B_{out}\right)\right) \qquad \text{(S.17)}$$

where the hemi-side function $\Phi(\ldots)$ describes the effective force exerted in the $z$ direction. Figure 4 E-G and SI Figure 4 C-D show the numerical results. In the simulation, we used $l = 3$, $r_0 = 1$, $\lambda_{Din0} = 100$, $\lambda_{Din1} = 1$, $\lambda_{Dout} = 0.1$, $B_{in} = 11$, $B_{out} = 0.65$, $C_0 = 20$, $\sigma = 0.1$, and $H_0 = 4$. For the effective force, we considered the dermal resistance and used Equations (S.15) and (S.17) to compute the force.

### References


1 Moore-Smith, L. D., Isayeva, T., Lee, J. H., Frost, A. & Ponnazhagan, S. Silencing of TGF-beta1 in tumor cells impacts MMP-9 in tumor microenvironment. *Sci Rep* **7**, 8678, doi:10.1038/s41598-017-09062-y

8678

10.1038/s41598-017-09062-y [pii]

9062 [pii] (2017).

2 Kobayashi, T. *et al.* Matrix metalloproteinase-9 activates TGF-beta and stimulates fibroblast contraction of collagen gels. *Am J Physiol Lung Cell Mol Physiol* **306**, L1006-1015, doi:10.1152/ajplung.00015.2014

ajplung.00015.2014 [pii]

L-00015-2014 [pii] (2014).

3 Strzyz, P. Forcing through barriers. *Nature Reviews Molecular Cell Biology* **20**, 136-136, doi:10.1038/s41580-019-0104-8 (2019).

4 Sadd, M. H. in *Elasticity* Ch. 5, 103 (Academic Press, 2014).

5 Wierzbicki, T. in *Structural Mechanics* 1.3.1-1.3.2 (LibreTexts, 2024).

6 Guo, C. L. Self-Sustained Regulation or Self-Perpetuating Dysregulation: ROS-dependent HIF-YAP-Notch Signaling as a Double-Edged Sword on Stem Cell Physiology and Tumorigenesis. *Front Cell Dev Biol* **10**, 862791, doi:10.3389/fcell.2022.862791 (2022).

7 Aragona, M. *et al.* A mechanical checkpoint controls multicellular growth through YAP/TAZ regulation by actin-processing factors. *Cell* **154**, 1047-1059, doi:10.1016/j.cell.2013.07.042 (2013).

8 Trepat, X. *et al.* Universal physical responses to stretch in the living cell. *Nature* **447**, 592-595, doi:10.1038/nature05824 (2007).

9 Constantinou, I. & Bastounis, E. E. Cell-stretching devices: advances and challenges in biomedical research and live-cell imaging. *Trends Biotechnol* **41**, 939-950, doi:10.1016/j.tibtech.2022.12.009 (2023).

10 Pourati, J. *et al.* Is cytoskeletal tension a major determinant of cell deformability in adherent endothelial cells? *Am J Physiol* **274**, C1283-1289, doi:10.1152/ajpcell.1998.274.5.C1283 (1998).

11 Na, S. *et al.* Time-dependent changes in smooth muscle cell stiffness and focal adhesion area in response to cyclic equibiaxial stretch. *Ann Biomed Eng* **36**, 369-380, doi:10.1007/s10439-008-9438-7 (2008).

12 Gladilin, E. *et al.* TGFbeta-induced cytoskeletal remodeling mediates elevation of cell stiffness and invasiveness in NSCLC. *Sci Rep* **9**, 7667, doi:10.1038/s41598-019-43409-x (2019).

13 Szeto, S. G. *et al.* YAP/TAZ Are Mechanoregulators of TGF-beta-Smad Signaling and Renal Fibrogenesis. *J Am Soc Nephrol* **27**, 3117-3128, doi:10.1681/ASN.2015050499 (2016).

14 Kim, O. V. *et al.* Compression-induced structural and mechanical changes of fibrin-collagen composites. *Matrix Biol* **60-61**, 141-156, doi:10.1016/j.matbio.2016.10.007 (2017).
15 Tronci, G., Doyle, A., Russell, S. J. & Wood, D. J. Triple-helical collagen hydrogels via covalent aromatic functionalization with 1,3-Phenylenediacetic acid. *J Mater Chem B* **1**, 5478-5488, doi:10.1039/C3TB20218F (2013).
16 Vinciguerra, P. *et al.* Imaging mass spectrometry by matrix-assisted laser desorption/ionization and stress-strain measurements in iontophoresis transepithelial corneal collagen cross-linking. *Biomed Res Int* **2014**, 404587, doi:10.1155/2014/404587 (2014).
17 Wollensak, G., Spoerl, E. & Seiler, T. Stress-strain measurements of human and porcine corneas after riboflavin-ultraviolet-A-induced cross-linking. *J Cataract Refract Surg* **29**, 1780-1785, doi:10.1016/s0886-3350(03)00407-3 (2003).
