## Supplementary video legends for "Novel Tissue Mechanics-Guided Cellular Flows Enable the Evolution of Feather Follicles"

### Supplementary Videos

**Supplementary Video 1. Feather bud protrusion.** Cell tracking video of E7 chicken explant culture for 18h reveals cell migration pattern changed from horizontal to vertical during bud protrusion.

**Supplementary video 2. Feather follicle invagination.** Cell tracking video of E10+24h explant showing different cellular flows.

**Supplementary video 3. Dermal papillae formation.** Cell tracking video of E17 flight feather to reveal differential cellular flows: (green) DP forming cells migrate downwards. (red) Epidermal tongue cells migrate downwards while distal feather epidermal cells migrate upwards. (yellow) Feather elongating dermal cells migrate distally.

**Supplementary video 4. Scutate scale formation.** Cell tracking video of E10+24h quail scale showing limited dermal cell migration during scale formation

**Supplementary video 5. Scale to feather conversion.** Cell tracking video of a newly induced feather bud on top of the scutate scale.
